# VLK drives extracellular phosphorylation of EphB2 to govern the EphB2-NMDAR interaction and injury-induced pain

**DOI:** 10.1101/2024.03.18.585314

**Authors:** Kolluru D. Srikanth, Hajira Elahi, Praveen Chander, Halley R. Washburn, Shayne Hassler, Juliet M. Mwirigi, Moeno Kume, Jessica Loucks, Rohita Arjarapu, Rachel Hodge, Stephanie I. Shiers, Ishwarya Sankaranarayanan, Hediye Erdjument-Bromage, Thomas A. Neubert, Zachary T. Campbell, Raehum Paik, Theodore J. Price, Matthew B. Dalva

## Abstract

Phosphorylation of hundreds of protein extracellular domains is mediated by two kinase families, yet the significance of these kinases is underexplored. Here, we find that the presynaptic release of the tyrosine directed-ectokinase, Vertebrate Lonesome Kinase (VLK/Pkdcc), is necessary and sufficient for the direct extracellular interaction between EphB2 and GluN1 at synapses, for phosphorylation of the ectodomain of EphB2, and for injury-induced pain. *Pkdcc* is an essential gene in the nervous system, and VLK is found in synaptic vesicles, and is released from neurons in a SNARE-dependent fashion. VLK is expressed by nociceptive sensory neurons where presynaptic sensory neuron-specific knockout renders mice impervious to post-surgical pain, without changing proprioception. VLK defines an extracellular mechanism that regulates protein-protein interaction and non-opioid-dependent pain in response to injury.

**One-Sentence Summary:** Synaptic protein-protein interactions and pain are regulated by the presynaptic release of the extracellular kinase VLK in the spinal cord.

---

Mass spectrometry analysis of the extracellular domains of proteins reveals hundreds of extracellular phosphorylation events on specific serine, threonine, and tyrosine residues (*1*). Phosphorylation of these residues is thought to be mediated by two families of kinases, but the functional significance of extracellular-directed kinases in the nervous system is unknown. Notably, biallelic loss-of-function mutations in the serine/threonine directed FAM20C kinase results in Raine syndrome (*2*), while the tyrosine directed Vertebrate Lonesome Kinase (VLK) is an essential gene in the mouse (*3*). These data indicate that further exploration of the function of these proteins will likely shed light into an underappreciated area of biology: the regulation of synaptic plasticity and behavior through extracellular phosphorylation.

One example of the importance of extracellular phosphorylation is the EphB-NMDAR interaction that occurs at excitatory synapses throughout the brain and spinal cord (*4, 5*). The EphB-NMDAR interaction is induced by the phosphorylation of a highly conserved extracellular tyrosine residue, Y504 in the Fibronectin-type III (FNIII) domain of EphB2 (*6*). The direct extracellular interaction, induced by Y504 phosphorylation, between EphBs and the NMDAR increases NMDA-type glutamate receptor channel open time and is required for the retention and clustering of normal numbers of NMDAR at synaptic sites, while disruption of the EphB-NMDAR interaction results in loss of NMDARs from synaptic sites (*7–9*). Extracellular phosphorylation of EphB2 at Y504 promotes EphB2’s interaction with the N-terminal domain (NTD) of GluN1 via a charge-mediated mechanism that regulates the mobility of GluN1 at synaptic sites (*5*). Phosphorylation of EphB2 at Y504 also promotes NMDAR-dependent pain in mice, suggesting an important physiological role for this extracellular phosphorylation and potentially providing new non-opioid pain therapeutic targets (*6*).

## Ectokinase VLK secretion is ephrin dependent and is sufficient and necessary for EphB2-NMDAR interaction

We hypothesized that an ectokinase might be required to induce the EphB2-NMDAR interaction by specifically phosphorylating extracellular tyrosine residues on EphB2 (*6*). To test this hypothesis, we assessed the ability of the VLK family of tyrosine-directed ectokinases to stimulate the EphB2-NMDAR interaction in cell lines, primary cortical neurons, and at spinal cord synapses from mouse and human tissue. Phosphorylation of Y504 on EphB2 occurs in the extracellular space, requires a protein-kinase that is secreted from neurons and ephrin-B2 stimulation of neurons releases a soluble, protein-based activity, that phosphorylates the EphB2 ectodomain at Y504 in an ATP-dependent manner (*6*). Therefore, we sought a protein kinase that satisfied two conditions: 1) it can induce the EphB2-NMDAR interaction, and 2) it is secreted from neurons in an ephrin-B2-dependent manner. To test whether any of the VLK-related tyrosine kinases (VLK, DIA1, DIA1R, FAM69A, FAM69B, and FAM69C) can induce the EphB2-NMDAR extracellular interaction, each of these six kinases (FLAG-tagged) was co-expressed with GluN1, GluN2B, and EphB2 in HEK293T cells. GluN1 co-immunoprecipitated with EphB2 from protein lysates only when either VLK or Fam69A was cotransfected (Fig. 1A), indicating that, of the six kinases only two, VLK and FAM69A, satisfy the first condition by inducing the EphB-NMDAR interaction.

**Fig. 1.**
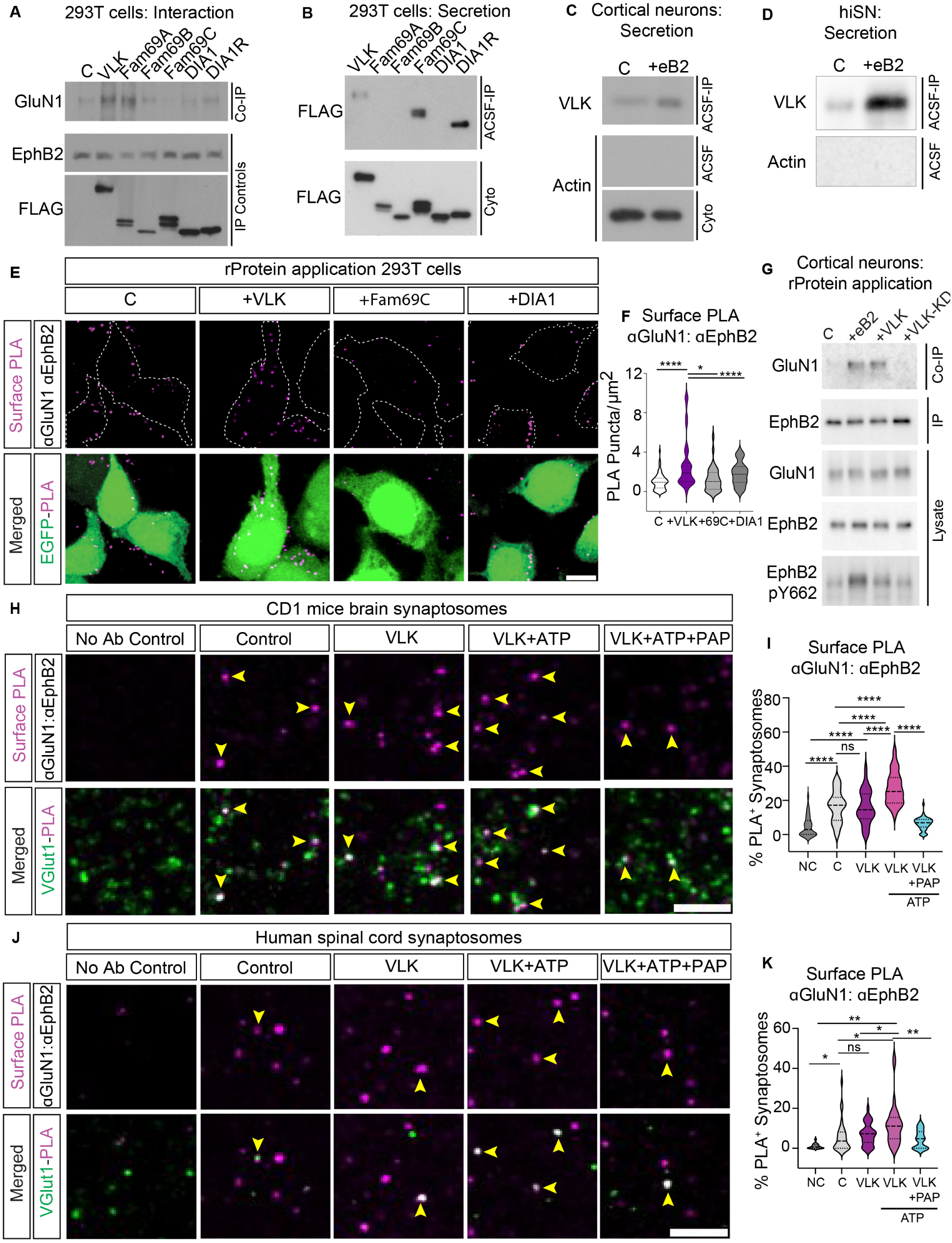
VLK is sufficient to induce the EphB2-NMDAR interaction. **(A)** Immunoblotting for co-immunoprecipitated GluN1 with EphB2 in stable VLK knockdown HEK293T cells (Co-IP, top). Control (C) cells were transfected with GluN1, GluN2B, and EphB2. Cells in other experimental conditions were transfected with VLK or each indicated VLK-related kinase FAM69A, FAM69B, FAM69C, DIA1, or DIA1R along with EphB2, GluN1 and GluN2B. Levels of EphB2 immunoprecipitated from cell lysates are shown by immunoblotting for EphB2 (IP Controls, middle). Extracellular kinases expression levels in cell lysates are shown by immunoblotting for FLAG after FLAG immunoprecipitation (FLAG, IP Controls, bottom). **(B)** Immunoblotting to detect the expression of secreted kinases in cell-conditioned ACSF (45min) from HEK293T cells transfected with kinases described above (ACSF-IP, top). Cell-conditioned ACSF and cell lysates (Cyto, bottom) were immunoprecipitated with a monoclonal anti-FLAG antibody and immunoblotted with a polyclonal anti-FLAG antibody. **(C)** Untransfected cultured cortical neurons (DIV7-8) treated with clustered EphrinB2 (+eB2) or with its respective control (C) reagents in ACSF for 45min. Phospho-**t**yrosine specific antibody pY99 was used to immunoprecipitate conditioned ACSF and immunoblotted with a custom-generated VLK antibody (ACSF-IP, top). Conditioned ACSF (ACSF, middle) and cell lysates (Cyto, bottom) were probed with beta-actin as a control to show that the conditioned media was free of any intracellular protein. **(D)** iPSC-derived human sensory neurons (hiSNs) cultured for four weeks and subject to control or clustered eB2 treatment as above. Experimental conditions for immunoprecipitation and immunoblotting as in (C) above. **(E)** Representative images of PLA of EphB2-NMDAR after treatment with soluble recombinant kinases. HEK293T cells were transfected with GluN1, GluN2B, EphB2, and EGFP. Cells were treated with active recombinant VLK, DIA1, or FAM69C proteins (as indicated) for 45 min before fixation. The upper panels show PLA signal alone (magenta). The lower panels are the merged images with EGFP in green. Scale bar is 10µm. **(F)** Quantification of the PLA puncta number. PLA puncta numbers were quantified by counting the puncta per 100 µm^2^ in EGFP^+^ cells and normalized to the Control. (****p=<0.0001, *p=0.038 One way ANOVA followed by Tukey’s; Control n = 70 fields from 7 repeats ; VLK n = 70 fields from 7 repeats; DIA1 n = 40 fields from 4 repeats; FAM69C n = 40 fields from 4 repeats ). **(G)** Untransfected cultured cortical neurons (DIV7-8) treated with control reagents (C), clustered EphrinB2 (+eB2), recombinant VLK (+VLK) or recombinant kinase dead VLK (+VLK-KD) for 45min. EphB2 was immunoprecipitated from the cell lysates, and co-immunoprecipitated GluN1 was immunoblotted (Co-IP, top). A fraction of the same sample was immunoblotted for EphB2 (EphB2, IP). Neuronal lysates were probed as lysate controls indicated for EphB2 and GluN1, and the intracellular phosphorylation of EphB2 was probed using EphB2 pY662. **(H)** Representative images of PLA of EphB2-NMDAR in CD1 mouse brain synaptosomes after treatment with recombinant VLK kinase with or without ATP or with ATP along with VLK with PAP phosphatase (as indicated). The upper panels show the PLA signal alone (magenta). The lower panels are merged images of synaptosomes immunostained for a presynaptic marker vGlut1 (green). The yellow arrows indicate colocalized puncta of presynaptic marker (vGlut1) and EphB2-NMDAR PLA puncta. The no antibody control condition was performed without primary antibody for EphB2 and GluN1. **S**cale bar is 10 µm. **(I)** Quantification of the percentage of vGlut1^+^ synaptosomes that colocalize with PLA puncta. (n= 40 fields from 4 different mice, One Way ANOVA followed by Tukey’s test ****p<0.0001). **(J)** Representative images of PLA of EphB2-NMDAR in Human Spinal cord synaptosomes. The yellow arrows indicate colocalized puncta of presynaptic marker (vGlut1) and EphB2-NMDAR PLA puncta. **S**cale bar is 10 µm. **(K)** Quantification of the percentage of vGlut1^+^ synaptosomes that colocalize with PLA puncta (n= 30, fields from 3 different human samples, One Way ANOVA followed by Tukey’s test ,*p=0.04,0.04,**p=0.006, 0.0067, )

To determine which of the kinases are secreted, each of the six kinases were overexpressed in HEK293T cells. After 24h, the culture media was replaced with a physiological salt solution (artificial cerebrospinal fluid; ACSF) for 45 minutes. FLAG immunoprecipitation from the conditioned ACSF (cACSF) revealed that only VLK, FAM69C, and DIA1R were constitutively secreted from HEK293T cells (Fig. 1B). Thus, only VLK is secreted and able to induce the EphB-NMDAR interaction when expressed in HEK293T cells.

To determine if endogenous VLK satisfies condition two and is secreted from neurons in an ephrin-B dependent manner, cultured rat cortical neurons at day *in vitro* seven (DIV7) and iPSC differentiated human sensory neurons (iSNs, Antomic), were treated with either control reagents (C) or activated ephrin-B2 (eB2) for 45 minutes in ACSF. The secreted tyrosine-phosphorylated proteins were immunoprecipitated from the cACSF and subsequent probing for VLK revealed that endogenous VLK was secreted by both rat cortical neurons and human iSNs (Fig. 1, C and D). Similarly, in cultured rat cortical neurons overexpressing VLK tagged with mCherry (VLK-mCherry), VLK was secreted upon treatment with activated ephrin-B2 (fig. S1A). Thus, in contrast to the constitutive secretion of VLK in HEK293T cells, VLK secretion from rat cortical neurons and hiSNs was regulated and elicited upon ephrin-B2 stimulation. In addition, ephrin-B2 stimulation selectively upregulated VLK mRNA (*PKDCC*) levels in rat cortical neurons and hiSNs (fig. S1, B-D). These data indicate that VLK can be secreted into the media by stimulation of neurons with ephrin-B2, that ephrin-B2 stimulation induces VLK expression, and that these effects are found in both rat and human neurons.

To test whether extracellular VLK is sufficient to induce the EphB2-NMDAR interaction, HEK293T cells were transfected with EphB2, GluN1, GluN2B, and EGFP. Transfected cells were treated with either control reagents, activated ephrin-B2, or purified recombinant (r) rVLK, rFAM69C, or rDIA1. Proximity ligation assay (PLA) was performed on unpermeablized transfected cells to determine whether the addition of recombinant kinases induced the EphB2-NMDAR interaction on the cell surface (*4*). Ephrin-B2 treatment robustly induced the EphB2-NMDAR interaction (fig. S1, E and F). Neither rFAM69C nor rDIA1 increased the EphB-NMDAR interaction compared to the control. In contrast, the rVLK application induced an EphB2-NMDAR interaction similar to the levels found with ephrin-B2 treatment (Fig. 1, E and F). Moreover, a kinase-dead mutant of VLK (rVLK-KD; fig. S1, E and F) failed to increase PLA indicating that VLK-dependent induction of the EphB2-NMDAR interaction requires VLK kinase activity.

To determine whether rVLK might induce the EphB2-NMDAR interaction in neurons, untransfected rat cortical neurons were treated with activated ephrin-B2, rVLK, or rVLK-KD. rVLK, but not rVLK-KD induced the endogenous EphB2-NMDAR interaction (Fig. 1G). To test whether EphB2 kinase activity is required for the EphB2-NMDAR interaction, activation of the EphB2 kinase was assayed using a phospho-specific EphB2 antibody (pY662, Fig. 1G bottom blot). Treatment of these untransfected neurons with rVLK induced the EphB2-NMDAR interaction without activation of the EphB2 kinase, suggesting that rVLK is sufficient to induce EphB2-NMDAR interaction even in the absence of ephrinB2 or induction of EphB2 kinase activity.

To test whether VLK might induce the EphB2-NMDAR interaction at synapses, we performed PLA for EphB2 and GluN1 in synaptosomes purified from mouse brain (Fig. 1, H and I) and human spinal cord from male and female donors (Fig. 1, J and K). Synaptosomes were treated with control reagents, rVLK, or rVLK with ATP. Addition of rVLK and ATP significantly increased the percentage of synaptosomes with EphB2 and GluN1 PLA puncta (Fig. 1, J-I) (*4*), suggesting that rVLK can induce the EphB2-NMDAR interaction at mouse cortical and human spinal cord synapses.

There are many phosphatases in the extracellular space (*10, 11*), suggesting that regulation of phosphorylation in this compartment may be biologically important. Consistent with this model, the extracellular phosphatase Prostatic acid phosphatase (PAP) is a negative regulator of pain in the spinal cord (*12*). Given the link between the EphB2-NMDAR interaction, pain, and extracellular phosphorylation, we asked whether PAP treatment might negatively regulate the EphB2-NMDAR interaction at mouse brain and human spinal cord synaptosomes. Remarkably, PAP blocked VLK-dependent increases in PLA puncta density in synaptosomes (Fig. 1, H-K). Together these data indicate that VLK is sufficient to induce the EphB2-NMDAR interaction and is likely the kinase mediating the extracellular phosphorylation of EphB2.

## Mechanism of VLK action on EphB2

Extracellular phosphorylation of EphB2 at Y504 imparts a negative charge on the FNIII domain of EphB2, which interacts with a positively charged pocket in the N-terminal hinge region of GluN1 to mediate EphB2-NMDAR interaction (*4, 6*). To determine whether rVLK phosphorylates the extracellular domain of EphB2, we conducted an *in vitro* cell-free kinase assay (Fig. 2A). Addition of rVLK and ATP but not rVLK-KD and ATP resulted in the robust phosphorylation of the EphB2 ectodomain (Fig. 2B). Neither of the secreted ectokinases, rFAM69C nor rDIA1 induced phosphorylation of the EphB2 ectodomain (Fig. 2C), suggesting that the extracellular domain of EphB2 is selectively phosphorylated by rVLK.

**Fig. 2.**
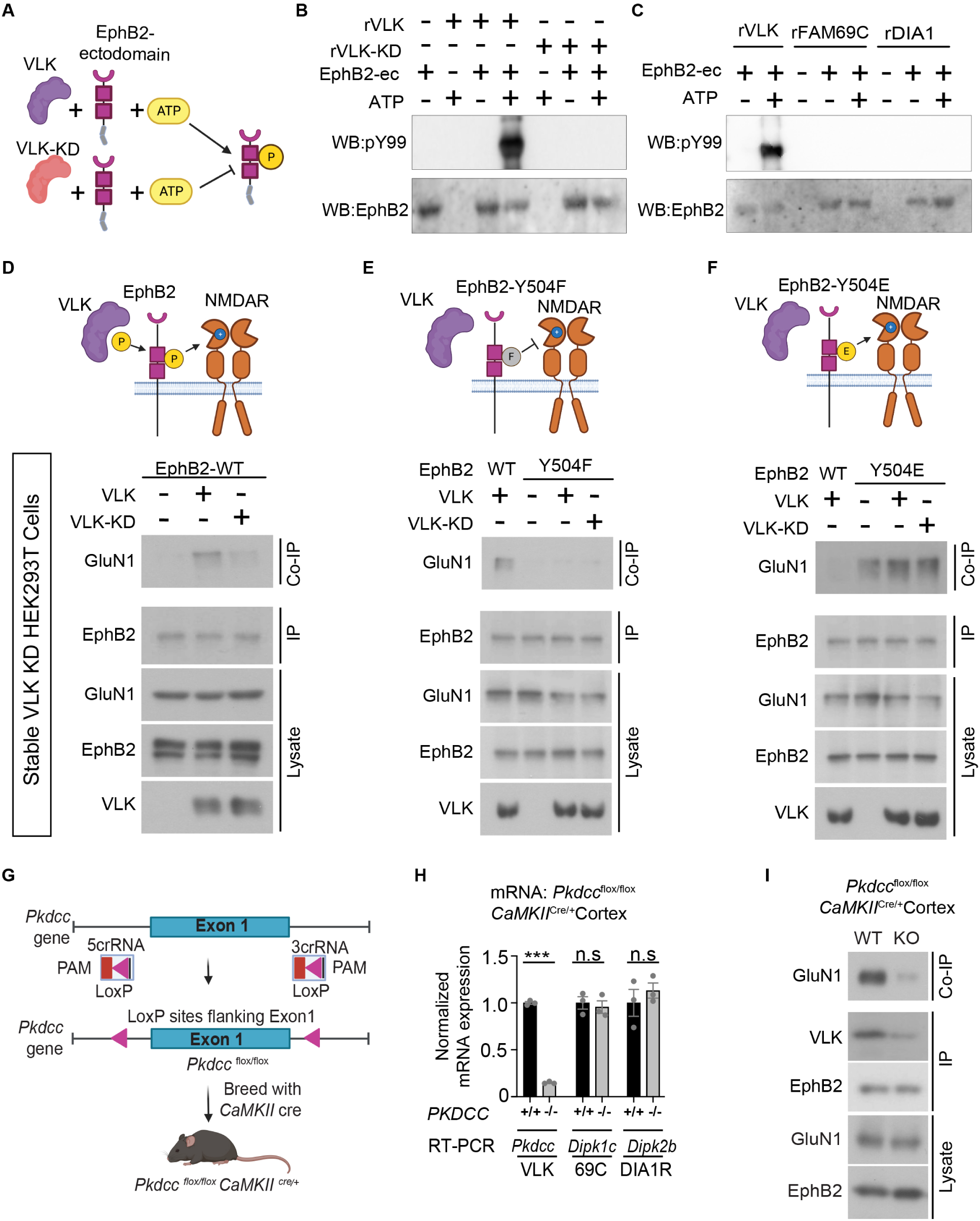
VLK extracellularly phosphorylates EphB2 and is necessary for the EphB2-NMDAR interaction. **(A)** Experimental approach to test *in vitro* EphB2 extracellular domain (EphB2-ectodomain) phosphorylation (yellow circle P) by active recombinant VLK (VLK, purple) or kinase-dead VLK (VLK-KD, orange). **(B)** *In vitro* kinase assay to test the tyrosine phosphorylation of EphB2 extracellular domain (EphB2-ec) by kinase active recombinant VLK (rVLK) or kinase-dead recombinant VLK (rVLK-KD) in ACSF, and MnCl_2_. The extracellular domain of EphB2 fused to the Fc fragment of immunoglobulin G (200ng) was incubated at 37°C for 30min with rVLK or rVLK-KD (100ng), either with or without ATP in MgAc (1mM) as indicated. EphB2-Fc was precipitated with Protein G agarose beads and probed with a phospho-tyrosine antibody pY99 and EphB2. **(C)** *In vitro* kinase assay to test the tyrosine phosphorylation of EphB2 extracellular domain (EphB2-ec) by kinase active recombinant FAM69C (rFAM69C) and Dia1 (rDia1). The experiment was set up as described above. rVLK has been included as a positive control for comparison. Immunoblots were probed as indicated. **(D-F)** Model illustrating VLK (purple) dependent extracellular phosphorylation (yellow circle P) at wildtype EphB2 tyrosine residue Y504 introduces a negative charge on EphB2 that is necessary for its interaction with the positively charged on the N-terminal domain of the NMDAR**(D)**. VLK cannot induce EphB-NMDAR interaction in the presence of a neutrally charged phospho-null mutant Y504F (grey circle F) in **(E),** and the interaction is constitutive in the presence of the negatively charged phosphor-mimetic mutant Y504E in the absence of VLK in **(F).** HEK 293T cells with stable VLK knockdown were transfected with GluN1, GluN2B, and either VLK or kinase-dead VLK (VLK-KD), and with wild type EphB2 in **(D)**, EphB2 Y504F in **(E)** or EphB2 Y504E in **(F)**. The cell lysates were immunoprecipitated for EphB2 and immunoblotted to detect co-immunoprecipitated GluN1. A fraction of the same EphB2 immunoprecipitated sample was immunoblotted for EphB2 (EphB2, IP), and lysate controls were probed as indicated. VLK blot is generated as described in Fig. 1B. **(G)** A schematic illustrating the conditional knockout of the *Pkdcc* gene that encodes for the VLK protein. *Pkdcc* floxed mice harboring loxP sites flanking the Exon 1 of the *Pkdcc* gene were bred with *CaMKII*-*Cre* to generate an excitatory cell-specific knockout *Pkdcc*^flox/flox^ *CaMKII*^Cre/+^ mice. **(H)** Real-Time (RT) PCR based quantification of relative mRNA levels of *Pkdcc* (VLK), *Dipk1c* (FAM69C), and *Dipk2b* (DIA1R) from the cortices of *Pkdcc*^flox/flox^ *CaMKII*^Cre/+^ mice (-/-, grey bars) compared to control mice cortices (+/+, black bars). (***p<0.0001, unpaired t-test; n=3). **(I)** Immunoblotting for co-immunoprecipitated GluN1 with EphB2 from cortical lysates of *Pkdcc*^flox/flox^ *CaMKII*^Cre/+^ (KO, top), compared to wild type (WT, top) littermate controls. A fraction of the same EphB2 immunoprecipitated sample was immunoblotted for EphB2 (EphB2, IP). pY99 immunoprecipitation from cortical lysate was immunoblotted for co-immunoprecipitated VLK (VLK, IP). Cortical lysates were probed as indicated for EphB2 and GluN1 (Lysate).

VLK can phosphorylate the ectodomain of EphB2, and a negative charge at Y504 on EphB2 is necessary and sufficient for the EphB2-NMDAR interaction (*6*). To determine whether VLK acts to induce the EphB2-NMDAR interaction by phosphorylating Y504, stable VLK knockdown HEK293T cells (fig. S2, A and B), which lacked constitutive EphB2-NMDAR interaction (Fig. 2D), were transfected with GluN1, GluN2B, and either wild type EphB2 (Fig. 2D; EphB2-WT), an EphB2 phospho-null mutant (Non-interacting mutant, Fig. 2E ; Y504F), or an EphB2 phospho-mimetic mutant (Constitutively interacting mutant, Fig. 2F; Y504E).

Consistent with the requirement for VLK kinase activity, the EphB2-NMDAR interaction was rescued by wild type VLK expression (Fig. 2D; +VLK ) but not by expression of the VLK-KD mutant (Fig. 2D; +VLK-KD ). Similarly, expression of VLK failed to induce the EphB2-NMDAR interaction in cells expressing EphB2-Y504F that cannot undergo phosphorylation at Y504 (Fig. 2E) and the EphB2-NMDAR interaction was constitutive in cells transfected with negatively charged constitutively interacting mutant EphB2-Y504E (Fig. 2F). These data indicate that VLK induces the EphB2-NMDAR interaction by phosphorylating Y504 on EphB2.

VLK is secreted by neurons and can phosphorylate the extracellular domain of EphB2. To test whether VLK is necessary for the EphB2-NMDAR interaction *in vitro* and *in vivo*, we engineered an inducible knockout of VLK mouse model by inserting lox-P sites around the first exon of *Pkdcc* (Fig. 2G). A neuron-specific VLK knockout mouse line was generated by crossing *Pkdcc*^flox/flox^ mice crossed with a *Nestin*-CRE line. However, no *Pkdcc* knockout mice were born from this line (0/106, 8 litters), suggesting that VLK is required in the nervous system development and viability. Therefore, we generated a forebrain excitatory neuron-specific knockout using the *CaMKII*-CRE line (Fig. 2G). *Pkdcc*^-/-^/*CaMKII*^Cre/+^ mice were viable and had a significant reduction of *Pkdcc* mRNA levels with no change in mRNA levels of the two other VLK-family secreted kinases, FAM69C (*Dipk1c*) or DIA1R (*Dipk2b;* Fig. 2H) in P35 mouse cortex. Consistent with the necessity of VLK for the EphB2-NMDAR interaction, GluN1 co-IP with EphB2 was reduced in cortex of *Pkdcc*^-/-^/*CaMKII*^Cre/+^ mice (Fig. 2I, *in vitro* data fig. S2C). These results show that VLK is necessary for the EphB2-NMDAR interaction.

## Presynaptic VLK regulates postsynaptic EphB2-NMDAR interaction

Ephrin-B activation of EphB2 results in the secretion of VLK from neurons. To determine the mechanism that mediates the release of VLK, we performed subcellular fractionation from the mouse forebrain to determine VLK subcellular localization. Surprisingly, VLK co-fractionated with clear core synaptic vesicles (Fig. 3A and fig. S3A) and not with dense core vesicles (fig. S3A). The presence of VLK protein in the synaptic vesicle fraction in mouse forebrain was confirmed using a targeted mass spectrometry analysis (fig. S3, B-E). These data suggest a model where VLK is positioned to be secreted from the presynaptic terminal to induce the postsynaptic interaction between EphB2 and the NMDAR.

**Fig. 3.**
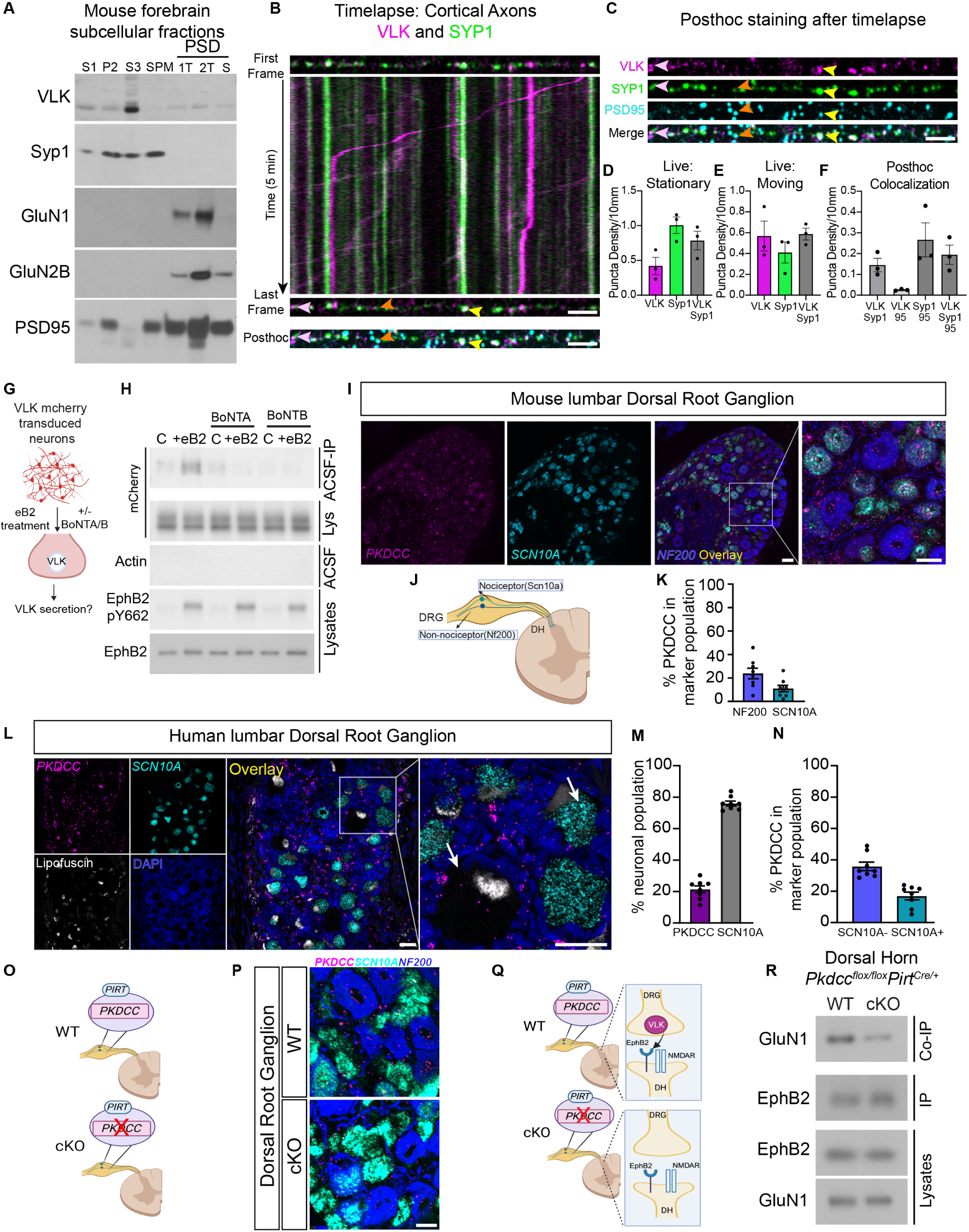
VLK is enriched in the presynaptic terminal and presynaptic VLK KO downregulates postsynaptic EphB-NMDAR interaction. **(A)** Western blots of PSD purification fractions prepared from WT CD1 mouse brain show VLK is enriched in crude vesicle fraction (S3). Gels were loaded with non-synaptic (S1), crude synaptosomal (P2), crude synaptic vesicle (S3), synaptic plasma membrane (SPM), and post-synaptic density (PSD) fractions. PSD fractions (from left to right) are least insoluble first triton extraction (1T), second triton extraction (2T), and most insoluble sarcosyl extraction (S). Blots were probed with VLK, Syp1, GluN1, GluN2B, and PSD-95 (as indicated). **(B)** The First and Last frame of the same axon segment from cultured rat cortical neurons (DIV21-23) transfected with VLK-mCherry (VLK, magenta), Synaptophysin1-EGFP (Syp1, green). mTurquoise was transfected as a cell fill (omitted for clarity). Kymograph generated from the indicated axon showing time (Y axis) vs distance (X axis). The same axon segment was fixed after live imaging and stained for PSD-95 (Posthoc, cyan). Scale bar is 5µm. Yellow arrowhead shows an example of colocalized VLK and Syp1 that remain stationary during live imaging and colocalize with PSD95. Pink arrowhead shows an example of colocalized VLK and Syp1 that remain stationary during live imaging but do not colocalize with PSD95. Orange arrowhead shows an example of Syp1 that remains stationary and colocalized with PSD95. **(C)** The individual channels for VLK, Syp1, PSD95 and Merged are shown for the same axon segment as (B) above. Scale bar is 5µm **(D-E)** Graph depicting the density of stable and moving VLK (magenta), Syp1 (green) and VLK+Syp1 (gray). **(F)** Graph depicting the puncta density of colocalized puncta after posthoc staining with PSD95 (95) VLK and Syp1 that remained stationary during live imaging and were maintained after fixation were quantified. **(G)** Schematic illustrating the experiment to test if ephrin B2 (eB2) stimulated VLK secretion from neurons can be blocked by botulinum toxins, A or B (BoNT A/B). **(H)** Cultured cortical neurons were transduced with lentivirus expressing VLK-mCherry at DIV3 and assayed at DIV7-8. Neurons were pretreated with BoNT A or B for 60min followed by treatment with clustered ephrinB2-(+eB2) or control (C) reagents with or without toxins in ACSF for 45-60min. Cell-conditioned ACSF were immunoprecipitated with RFP Trap agarose and immunoblotted for mCherry (top blot, ACSF IP). Cell-conditioned ACSF were probed for beta-actin to show no intracellular protein leaked into the ACSF (Actin, Sup). Cell lysates were probed for mCherry, EphB2 and EphB2 pY662 (as indicated). **(I)** RNAscope in situ hybridization for *Pkdcc* (Magenta) in mouse lumbar Dorsal Root Ganglion (DRG) , in *Scn10a* nociceptors (cyan) and Merged overlay with Nf200 mechanoreceptor cells (blue). Scale bar is 50 µm. **(J)** Mouse Dorsal Root Ganglion (DRG) have both nociceptor and mechanoreceptor cells marked by Scn10a and Nf200 respectively. **(K)** Quantification of High *Pkdcc* expressing neurons (n=8, Nf200 Mean = 23.83, SEM ± 4. 490, Scn10a Mean = 10.95, SEM ± 2.740). **(L)** RNAscope in situ hybridization for *PKDCC* (Magenta) in human lumbar DRG, *SCN10**A*** (**cyan**), Lipofuscin (white), DAPI (blue) Scale bar is 50 µm. Merged overlay, scale bar is 50 µm. Inset from overlay showing *PKDCC* expression in neurons (white arrows) Scale bar is 50 µm. **(M)** Quantification of *PKDCC* expressing neurons (n=8, *PKDCC* Mean = 21. 41, SEM ± 2. 156; *SCN10A* Mean = 75.98, SEM ± 1.509). (**N)** Quantified percentage of *PKDCC* expression in nociceptor and non-nociceptor populations. (n=8, Non-nociceptor Mean = 35.82, SEM ± 2.706; Nociceptor Mean = 16.95, SEM ± 2. 443). **(O)** A model of the DRG from *Pkdcc* conditional knockout mice (cKO). *Pkdcc*-flox mice were crossed to *Pirt*-Cre line to generate *Pkdcc* sensory-neuron specific cKO mice **(P)** RNAscope in situ hybridization for *Pkdcc* to confirm *Pkdcc* deletion from sensory neurons in *Pkdcc* cKO mice. Scale bar is 20 µm **(Q)** Model of WT and cKO mice (as described above) detailing the experiment to test the effect of *Pkdcc* deletion in the presynaptic DRG sensory neurons on the EphB2-NMDAR interaction in the dorsal horn (DH) postsynaptic neurons. **(R)** Immunoblot showing levels of GluN1 co-immunoprecipitated with EphB2 in the DH of cKO mice compared to wildtype littermate controls (WT). A fraction of the same sample was immunoblotted for EphB2 (EphB2, IP). DH lysates were probed as indicated for EphB2 and GluN1.

If VLK functions presynaptically to induce the EphB2-NMDAR interaction by being released with synaptic vesicles, we expect that: 1) VLK should localize to synaptic sites and co-traffic in axons with markers for synaptic vesicles, 2) blocking synaptic vesicle fusion should block ephrin-B dependent secretion of VLK, and 3) knocking out VLK presynaptically should block the EphB2-NMDAR interaction in postsynaptic neurons. To determine whether VLK localizes to presynaptic terminals, we asked whether VLK is found in axons and traffics with synaptic vesicle markers. Timelapse imaging of live neurons co-expressing VLK-mCherry and Synaptophysin1-EGFP (Syp1-EGFP) revealed that both stable and moving VLK puncta colocalized with Synaptophysin1 (Fig. 3B, D-E). Consistent with fast axonal transport (*13*), the average velocity of VLK was 0.6 ±0.4 μm/sec (Fig. 3B and movie S1). *Post hoc* staining for PSD-95, VLK and Syp1 shows that VLK is preferentially localized to synaptic sites (Fig. 3, C and F). These data suggest that VLK traffics in axons with presynaptic proteins and is found presynaptically at many excitatory synapses.

Ephrin-B activation of EphB2 induces the secretion of VLK (Fig. 1, C and D). To test whether ephrin-B-induced VLK release is mediated by a SNARE-dependent mechanism, rat cortical neurons, transduced with VLK-mCherry lentivirus, were pretreated with either Botulinum toxin A or B (BoNT-A/B) to cleave SNAP-25 or synaptobrevin respectively and treated with ephrin-B2 for 45 minutes (Fig. 3G). Cleavage of the SNARE complex with either BoNT-A or BoNT-B blocked the release of VLK-mCherry and reduced levels of VLK in the media with and without ephrin-B2 stimulation of EphB kinase signaling but did not block ephrin-B2 dependent activation of EphB2 (Fig. 3H). This shows that VLK is released into the media by a mechanism that requires the SNARE complex. Thus, VLK copurifies and colocalizes with presynaptic vesicle makers, and secretion of VLK is abrogated by blocking synaptic transmission.

If presynaptic axonal terminals release VLK to induce the EphB2-NMDAR interaction in postsynaptic cells, removal of VLK from axons should block the EphB2-NMDAR interaction. Next, we sought to identify a circuit where we could isolate presynaptic terminals of projection neurons. One such circuit is formed by the dorsal root ganglion (DRG) neurons that project into the dorsal lamina of the spinal cord (*14*). The EphB2-NMDAR interaction is pathologically upregulated in the spinal cord by injury of peripheral tissues which drives activation of DRG neurons (*15*), suggesting that DRG neurons might express VLK. Consistent with this model, the VLK transcript, *Pkdcc* is detected in mouse DRG by single-cell RNAseq (fig. S4, A-B) and RNAscope *in situ* hybridization revealed *Pkdcc* mRNA expression in both mechanoreceptor (NF200*^+^*: 24%) and nociceptor (*Scn10a^+^*: 11%) populations of the DRG (Fig. 3, I-K and fig. S4C). The expression pattern of *Pkdcc* was similar in both female and male mice (fig. S4, D-H). RNAscope in human tissue obtained from male and female organ donors (table S1) found *PKDCC* expression in 21% of human DRG neurons (Fig. 3, L and M). The *PKDCC* positive neurons were small to medium-sized and had similar expression patterns in the two sexes (fig. S5, A and B), with *PKDCC* expressed in 17% of nociceptors and 36% of non-nociceptors (Fig. 3N). VLK protein expression in the human lumbar DRG was confirmed using the Somascan proteomic platform (fig. S5C). These data indicate that in both mouse and human DRG, expression of *PKDCC* is found in neurons associated with nociception that send axonal projections to the dorsal lamina of the spinal cord.

Based on the pattern of VLK transcript expression in the DRG, we directly tested whether VLK is required presynaptically to induce the EphB2-NMDAR interaction. By crossing the *Pirt*-CRE driver line with the *Pkdcc*^flox/flox^ mice (*16*), VLK was knocked out in all DRG sensory neurons (Fig. 3, O and P) without altering *Pkdcc* mRNA (VLK) levels in the spinal cord (fig. S5D). The dorsal horn region of the spinal cord was isolated, leaving only the DRG axon terminals and not the cell bodies from the *Pkdcc*^-/-^*Pirt^Cre/+^* knockout mice. In the VLK knockout spinal cord (VLKcKO), the co-immunoprecipitation of GluN1 with EphB2 from the dorsal horn was significantly decreased compared to wild type controls (Fig. 3, Q and R). These data indicate that loss of VLK from axons is sufficient to block the EphB2-NMDAR interaction and support a model in which axon terminals secrete VLK to regulate the ability of EphB2 and the NMDAR to interact directly via their extracellular domains. Accordingly, VLK is found in the presynaptic terminal, is released in a SNARE-dependent fashion from synapses, and functions presynaptically to induce the EphB2-NMDAR interaction, indicating that VLK is a synaptically released essential mediator of the EphB2-NMDAR interaction.

## VLK drives a pain-like state in mice dependent on kinase activity and the NMDAR

In acute and chronic pain states, NMDARs are activated by sustained nociceptive stimulation, and NMDAR activation mediates both short and long-term plasticity that enhances pain detection by amplifying ascending nociceptive neurotransmission to the brain (*17*). Upregulation of NMDAR function by induction of the EphB-NMDAR interaction via overexpression of a phosphomimetic mutant EphB2 receptor (Y504E) induces pain hypersensitivity in mice. Moreover, in models of surgical wound pain, the EphB-NMDAR interaction is increased (*6*). These data suggest that extracellular phosphorylation of EphBs may drive NMDAR-dependent hypersensitivity and pain plasticity. Therefore, we tested whether the ectokinase VLK might mediate pain plasticity *in vivo*.

To test whether VLK can induce pain-like behaviors, rVLK was injected intrathecally into naïve mice targeting the lumbar spinal cord and DRGs (Fig. 4A). Mice received rVLK or VLK family kinases, rFAM69C or rDIA1, or heat-denatured recombinant proteins. Effects on mechanical sensitivity were determined with von Frey filaments and spontaneous pain by the mouse grimace scale (*18*). Neither rDIA1 nor rFAM69C produced any changes in grimacing or mechanical sensitivity. In contrast, injection of rVLK resulted in a significant increase in mechanical sensitivity and grimace (Fig. 4, B and C). These effects were specific to VLK kinase activity, as heat-denatured rVLK or the rVLK-KD mutant did not cause any behavioral changes (Fig. 4, D and E). The extracellular phosphatase PAP is natively expressed by a population of nociceptors, negatively regulates pain hypersensitivity (*12*), and the addition of rPAP blocks VLK-dependent induction of the EphB2-NMDAR interaction in synaptosomes (Fig. 1, H-K). Therefore, we asked whether pre-injection of rPAP might block the induction of mechanical hypersensitivity by VLK (Fig. 4F). Consistent with this model, pretreatment with PAP blocked the effects of rVLK injection (Fig. 4, F and G). However, rPAP pretreatment did not affect grimace behaviors (Fig. 4H), suggesting that the PAP phosphatase cannot suppress all of the effects of rVLK-induced pain. Collectively, the effect of rPAP on rVLK-induced mechanical hypersensitivity suggests a model where endogenous phosphatase activity may limit the activity of VLK released from DRG neurons.

**Fig. 4.**
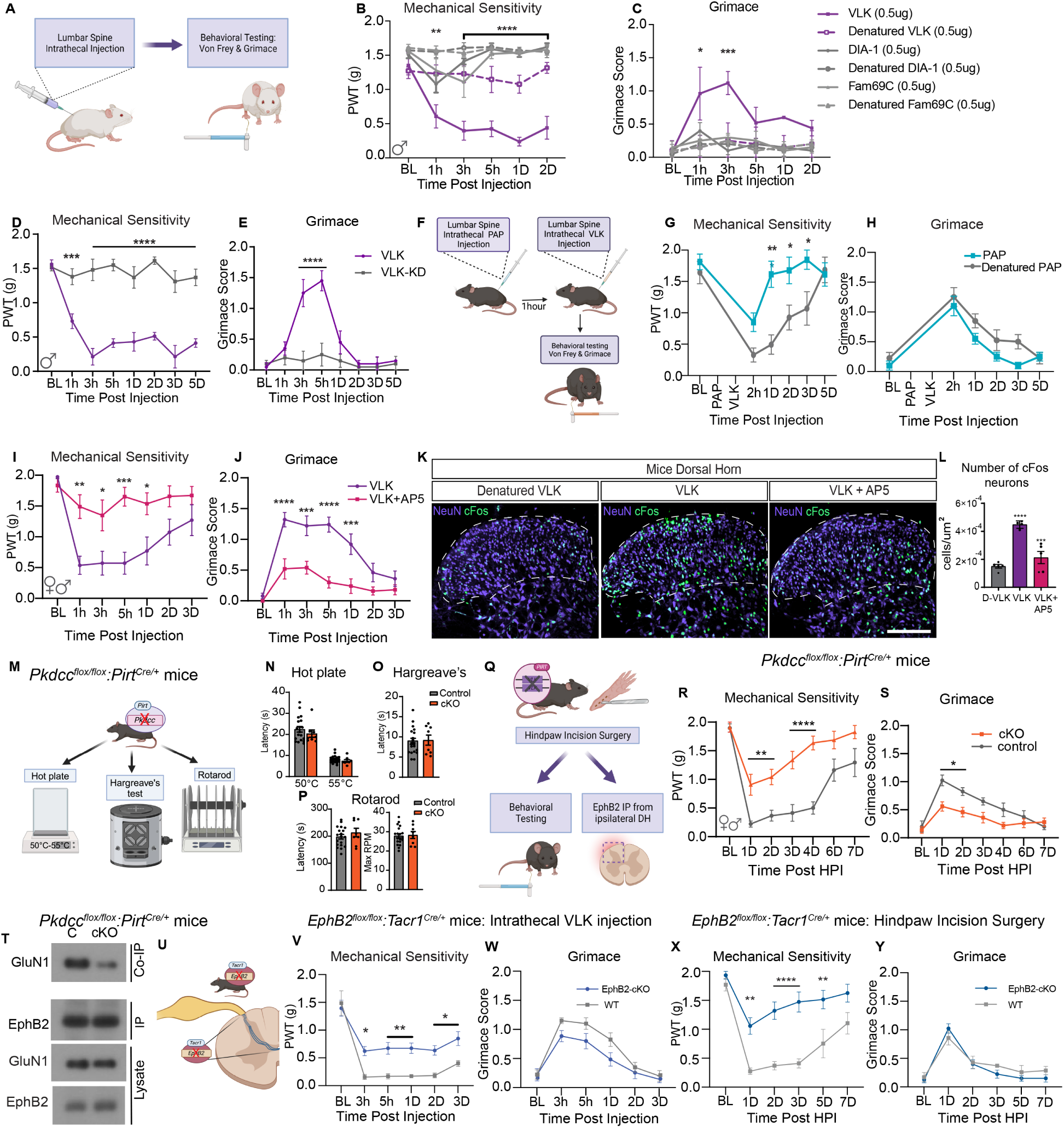
VLK induces a robust pain state dependent on NMDA receptor activation, and endogenous expression in DRG sensory neurons is necessary for post-operative pain. **(A)** Schematic illustration of intrathecal drug administration and measurement of pain-like behaviors. von-Frey testing is used as a measure of mechanical sensitivity and grimace scores depict levels of affective pain. **(B, C)** Three different secreted kinases and their denatured controls were administered to mice with 0.5μg of VLK, DIA-1 and FAM69C. Mice receiving VLK displayed a sharp drop in mechanical thresholds and increased grimacing. (n = 4, Two-way repeated measures ANOVA, F (5, 162) = 82. 33,**p=0.001, ****p<0.0001; F (5, 162) = 9.759, ***p<0.0001. post hoc Dunnett’s test). **(D, E)** Administration of a kinase-dead mutant of VLK (VLK-KD) abolishes pain-like behaviors induced by VLK. (n=5, Repeated measures ANOVA, F (2, 11) = 26. 19, ****p<0.0001; Two-way repeated measures ANOVA, F (2, 11) = 11.07, **p=0.0023. post hoc: Dunnett’s test). **(F)** Schematic of administration of the extracellular phosphatase PAP followed by VLK and measurement of pain-like behaviors. **(G, H)** Mice administered PAP followed by VLK showed significantly reduced mechanical sensitivity (n=8, Two-way repeated measures ANOVA, F (5, 70) = 18.96,***p=0.0003 Posthoc: Bonferroni) whereas PAP did not affect VLK-induced grimacing compared to denatured PAP control (n=8, Two-way repeated measures ANOVA, posthoc: Bonferroni) . **(I, J)** Co-administration of VLK with AP5 also blocked the VLK-induced pain phenotype. (n=8, Two-way repeated measures ANOVA F (1, 18) = 18.85, ***p=0.0004, F (1, 18) = 19.93, ****p=0.0003. post hoc: Bonferroni). **(K)** The lumbar spinal cord dorsal horn from mice administered Denatured VLK, VLK, or VLK + AP5 was immunohistochemically stained for NeuN and cFos markers. **(L)** Quantification of cFos^+^ cells showed VLK induced a significant increase in neuronal cFos expression which was blocked by AP5 antagonism of NMDARs or with the use of heat Denatured VLK (D-VLK). (One-way ANOVA with Bonferroni multiple comparisons ****p<0.0001, ***p<0.001 **(M)** Model depicting *Pkdcc*^flox/flox^ *Pirt*^Cre/+^ mice (cKO) and *Pkdcc*^flox/flox^ littermate controls (Control) were subjected to behavioral testing for normal sensorimotor development. (**N-P**) Paw withdrawal latencies for were measured on a Hot Plate in **(N)** and Hargreaves assay in (**O).** Rotarod testing was performed in **(P)**. No significant differences were observed between genotypes in the above tests. **(Q)** A model of the *Pkdcc* conditional knockout mice (cKO). *Pkdcc*-flox mice were crossed to *Pirt*-Cre line to generate *Pkdcc* sensory-neuron specific cKO mice. cKO and wild-type littermates were subjected to Hind-paw incision surgeries. **(R-S)** Behavioral data showing effects of sensory neuron-specific VLK knockout on postsurgical mechanical hypersensitivity and affective pain. (n = 9 CKO, n = 8 Littermate; Two-way repeated measures ANOVA F (1, 16) = 21. 38, ***p=0.0003; F (1, 16) = 9.902, **p=0.0062. posthoc: Bonferroni). **(T)** Immunoblot showing levels of GluN1 co-immunoprecipitated with EphB2 in the ipsilateral dorsal horn (DH) in cKO mice compared to wildtype littermate controls (C). A fraction of the same sample was immunoblotted for EphB2 (EphB2, IP). DH lysates were probed as indicated for EphB2 and GluN1. **(U)** Illustration depicting intrathecal injection of recombinant VLK in *EphB2*^flox/flox^ *Tacr1*^Cre/+^ mice (EphB2 cKO). (**V-W**) Measurement of pain-like behaviors using Von Frey **(V)** (n=7, Two-way repeated measures ANOVA, F (5, 65) = 2.641, **p=0.003, posthoc: Bonferroni) and Grimace **(W)** (n=7, p=NS, posthoc: Bonferroni). **(X-Y)** Behavioral data (Von-Frey and Grimace) of *EphB2*^flox/flox^ *Tacr1*^Cre/+^ mice (EphB2 cKO) following Hindpaw Incision surgery. Mechanical Sensitivity measured using Von Frey **(X)** (n = 8 CKO, n = 7 Littermate; Two-way repeated measures ANOVA F (5, 65) = 3.557, **p=0.0066, ****p<0.0001, posthoc: Bonferroni), Grimace **(Y)** (n = 8 CKO, n = 7 Littermate; Two-way repeated measures ANOVA p=ns, posthoc: Bonferroni).

rVLK induces the EphB2-NMDAR interaction, and the induction of the EphB2-NMDAR interaction increases the synaptic localization of the NMDAR, open time of the channel, and NMDAR-dependent calcium influx in neurons while also inducing mechanical pain hypersensitivity (*6, 7*). To determine whether rVLK might act to modulate pain behavior through the NMDAR, we blocked the NMDAR in the presence of rVLK. Intrathecal injection of naïve mice with rVLK and the NMDAR antagonist, AP5, prevented the rVLK-dependent induction of a pain-like phenotype (Fig. 4, I and J). Stimulation of synaptic activity associated with pain-like behavior induces expression of activity-dependent immediate early genes like *cFos* (*19*). To test whether *cFos* activity can be stimulated by VLK, mice received intrathecal injection of rVLK, rVLK with the NMDAR blocker AP5, or an rVLK-KD mutant protein. Ninety minutes following the injections, spinal cords were processed for immunostaining. *cFos* positive cells were observed in the superficial and deeper lamina of the dorsal horn where projection neurons that send ascending nociceptive signaling to the brain are found (Fig. 4K). The effect of active rVLK on *cFos* expression was significantly greater than with NMDAR blockade or with the treatment of the rVLK-KD (Fig. 4L). Accordingly, rVLK drives the induction of pain-like behaviors and increases NMDAR-dependent neuronal activity, linking VLK-dependent induction of the EphB2-NMDAR interaction directly to pain plasticity.

## Sensory neuronal VLK expression mediates surgical pain

rVLK is sufficient to induce pain-like behaviors. Extracellular phosphorylation of EphB2 is a key mechanism driving the EphB2-NMDAR interaction in the dorsal horn and mechanical pain hypersensitivity in the postsurgical pain model (*6*). rVLK administration is sufficient to produce pain-like behaviors in naïve mice and induce the EphB-NMDAR interaction, and VLK mRNA is expressed in DRG neurons. To test the necessity of VLK in the development of mouse postsurgical pain, *Pkdcc*/VLK was deleted in sensory neurons using *Pirt-Cre* mice (Fig. 4M). The *Pkdcc*^-/^*Pirt^Cre/+^* mice had no defects in hot plate or Hargreaves’ tests of thermal nociception and performed normally on the rotarod assay, showing that although rVLK can induce pain-like responses in mice, expression of *Pkdcc* is not required for normal sensory-motor behavior (Fig. 4, N-P).

To test whether VLK regulates injury-induced pain-like responses in mice, the response of *Pkdcc*^flox/flox^*Pirt^Cre/+^* mice to surgical wound pain was measured. Hind paw incision surgery was performed on VLK-cKO and VLK-WT littermates (Fig. 4Q). The control group showed an acute increase in mechanical hypersensitivity that took up to a week to return to baseline. Strikingly, the mechanical sensitivity was significantly attenuated in the VLK-cKO group and returned to baseline by day 4 (Fig. 4R). In addition to differences in mechanical sensitivity, the VLK-cKO group also showed less facial grimacing after surgery, indicating a decrease in spontaneous pain (Fig. 4S). These data indicate that VLK drives injury-induced mechanical hypersensitivity and spontaneous pain behaviors but is dispensable for normal sensory and sensory-motor activity.

VLK is expressed in DRG neurons and is required there for injury-induced pain signaling. To begin to determine whether this effect was caused by the selective expression of VLK in only DRG neurons, the expression pattern of *PKDCC* was determined in the dorsal horn of spinal cord of mice and humans. In mice *Pkdcc* mRNA expression was distributed broadly across dorsal horn neurons (41% of neurons), with substantial overlap (64%) with *Tacr1*^+^ neurons (fig.S6), a population of neurons that includes projection neurons of the spinothalamic tract involved in ascending pain signaling (*20, 21*). Similarly, the human organ-donor spinal cord showed expression of *PKDCC* in 60% of dorsal horn neurons, including large neurons in the outer lamina that are likely projection neurons (fig. S7). Thus, *Pkdcc* is expressed in both the spinal cord and DRG, suggesting that the decreased mechanical sensitivity in the DRG VLK cKO mice may be driven by the loss of presynaptic VLK, which in turn reduces the EphB2-NMDAR interaction and pain plasticity.

To determine whether the decreases in pain-like behavior in VLK sensory neuron-specific cKO mice might be related to loss of the VLK-dependent EphB2-NMDAR interaction in the dorsal horn of the spinal cord, we first validated that DRG-specific knockout of *Pkdcc*, had no effect on the levels of *Pkdcc* mRNA in the spinal cord (fig. S5D). Next, the ability of EphB2 and NMDAR to interact was determined by co-immunoprecipitation from *Pkdcc*^-/-^*Pirt^Cre/+^* mice and littermate controls following hind paw incision surgery. Twenty-four hours after surgery, *Pkdcc*^-/-^*Pirt^Cre/+^* and littermate controls dorsal horns had similar levels of expression of EphB2 and GluN1, but the EphB2-NMDAR interaction was lost in the dorsal horn of spinal cord of DRG specific *Pkdcc* knockout mice (Fig. 4T). These data indicate that VLK is necessary for injury-induced increases in the EphB2-NMDAR interaction and consistent with a model where the EphB2-NMDAR interaction serves as an underlying mechanism modulating NMDAR function in injury-induced pain signaling in DRG to spinal cord circuits.

Pain information is carried to the brain via *Tacr1*^+^ projection neurons, and postsynaptic EphBs are required for the EphB-NMDAR interaction. Analysis of RNA sequencing databases indicates that *EPHB2*, *GRIN1* (GluN1), *TACR1*, and *PKDCC* are co-expressed by excitatory dorsal horn neurons in human and mouse spinal cord, including in verified projection neurons in mice (*22–24*). To test whether VLK-driven pain-like behaviors require EphB2 expression in *Tacr1*^+^ spinal cord neurons, we generated an *EphbB2^flox/flox^ Tacr1*^Cre/+^ mouse (Fig. 4U;(*25*), EphB2-cKO). Although the *Tacr1*^Cre/+^ is only expressed in about 40% of the *Tacr1* population of projection neurons (*20*), intrathecal injection of VLK in *Tacr1 EphB2* knockouts, resulted in a significant decrease in mechanical hypersensitivity (Fig. 4V). Moreover, the surgical wound model failed to induce mechanical hypersensitivity in the *EphB2^-/-^ Tacr1*^Cre/+^ KO mice. These findings indicate that EphB2 expression on the *Tacr1*^+^ neurons of the spinal cord mediates the effects of VLK in producing mechanical hypersensitivity after injection and injury. However, in neither of these experiments were grimace levels affected by genotype (Fig. 4, W and Y). The lack of effect on grimacing may reflect that a different subset of ascending projection neurons carry information on acute pain behaviors, like grimacing, after injury in mice (*26*) or may be due to the small percentage of projection neurons targeted in the *Tacr1*-CRE line (*20*). Collectively, these results indicate that VLK expression in sensory afferents is necessary for the development of post-surgical pain and demonstrate that extracellular kinase activity is essential for the regulation of protein-protein interactions at synapses onto spinal neurons that express EphB2 to regulate pain-related behavioral responses to injury.

## Discussion

We describe a previously unappreciated role for the ectokinase VLK in protein-protein interaction, synaptic receptor localization, and injury-induced pain behaviors. Our data support a model in which the extracellular kinase VLK is released presynaptically, likely from small clear core vesicles, also known to co-release glutamate and ATP (*6, 27*), to regulate phosphorylation of EphB receptors on a specific extracellular residue, leading to clustering of NMDARs at synapses (*6*). The phosphorylation of EphB2 occurs on a specific, highly conserved tyrosine residue, found in Eph proteins across phylogeny and conserved in many other synaptic FNIII-containing proteins (*6*). Neuronal expression of sVLK is necessary for viability and essential for NMDAR-dependent injury-induced pain. Accordingly, extracellular phosphorylation may be important more broadly in synaptic plasticity and behavior due to the widespread requirement for NMDARs (*28, 29*). Our work creates a set of tools and a unique perspective on synaptic biology that will likely enable new insight into ectokinase function. This is likely important beyond VLK because mutations in other kinases in this family are linked to CNS diseases (*1, 30*).

The ability to respond to injury with amplified nociceptive signaling is critical for the survival of multicellular organisms with complex nervous systems (*17*). Augmented NMDAR activity at the DRG to spinal cord synapse (PNS to CNS) is a conserved mechanism in vertebrates (*17, 31*). However, therapeutic intervention at the NMDAR is limited by the strong CNS effects that arise when inhibiting this receptor (*32*). Our findings demonstrate a different approach to targeting the NMDAR complex in the context of pain, a strategy expected to leave normal sensory activity in place while decreasing the augmented activity that drives injury-induced pain amplification. Targeting the VLK/EphB2/NMDAR complex, which appears to be present in both mouse and human spinal cord synapses, in the spinal cord could provide a long-sought-after, non-opioid mechanism to manipulate NMDAR receptor-dependent pain without severe side effects that are found in current approaches like receptor antagonists. There will be limitations to this approach as indicated by the findings that, while elimination of VLK from sensory neurons or blocking NMDARs during rVLK treatment blocked both mechanical hypersensitivity and grimacing in mice, spinal manipulation of EphB2 gene expression in a small subset of projection neurons only affected mechanical hypersensitivity. These data highlight the importance of future work to understand VLK’s modality-specific functions, targeting of specific projection neuron subtypes, and effect on local spinal circuits in pain sensation and plasticity. Additional opportunities come from the conservation of VLK expression, release, and effects on the EphB-NMDAR interaction in mice and humans, highlighting the translational potential of this non-opioid-dependent approach.

## Acknowledgments

The authors thank Erin Vines, Peter Horton, Geoffrey Funk, Anna Cervantes and others at the Southwest Transplant Alliance for recovery of tissues from organ donors. The authors thank the organ donors and their families for their gift. We thank Dr Mark Zylka at University of North Carolina, Chapel Hill, for the gift of recombinant PAP. We thank Patrick Walsh and Vince Truong for their assistance with iSN work. We thank members of the Price and Dalva lab for advice.

## Author contributions

Conceptualization: TJP and MBD

Methodology: KDS, HE, PC, HRW, RP, ZC, TAN, TJP, MBD

Investigation: KDS, HE, PC, HRW, SH, MK, JMM, JL, RA, SIS, RP, IS, HEB

Visualization: KDS, HE, PC, HRW, HEB Funding acquisition: TJP, MBD

Project administration: TJP, MBD Supervision: TJP, MBD

Writing – original draft: KDS, HE, PC, HRW, TJP, MBD Writing – review & editing: KDS, HE, PC, HRW, TJP, MBD

### Data and materials availability

All data are available in the main text or the supplementary materials. Raw data files supporting the findings of the manuscript are available from the corresponding authors Theodore J. Price and Matthew B. Dalva

## References

1. V. S. Tagliabracci et al., A Single Kinase Generates the Majority of the Secreted Phosphoproteome. Cell 161, 1619–1632 (2015).

2. V. S. Tagliabracci et al., Secreted kinase phosphorylates extracellular proteins that regulate biomineralization. Science 336, 1150–1153 (2012).

3. Y. Imuta, N. Nishioka, H. Kiyonari, H. Sasaki, Short limbs, cleft palate, and delayed formation of flat proliferative chondrocytes in mice with targeted disruption of a putative protein kinase gene, Pkdcc (AW548124). Dev Dyn 238, 210–222 (2009).

4. H. R. Washburn, N. L. Xia, W. Zhou, Y. T. Mao, M. B. Dalva, Positive surface charge of GluN1 N-terminus mediates the direct interaction with EphB2 and NMDAR mobility. Nat Commun 11, 570 (2020).

5. H. R. Washburn, P. Chander, K. D. Srikanth, M. B. Dalva, Transsynaptic Signaling of Ephs in Synaptic Development, Plasticity, and Disease. Neuroscience 508, 137–152 (2023).

6. K. Hanamura et al., Extracellular phosphorylation of a receptor tyrosine kinase controls synaptic localization of NMDA receptors and regulates pathological pain. PLoS Biol 15, e2002457 (2017).

7. M. J. Nolt et al., EphB controls NMDA receptor function and synaptic targeting in a subunit-specific manner. J Neurosci 31, 5353–5364 (2011).

8. M. S. Kayser, A. C. McClelland, E. G. Hughes, M. B. Dalva, Intracellular and trans-synaptic regulation of glutamatergic synaptogenesis by EphB receptors. J Neurosci 26, 12152–12164 (2006).

9. M. B. Dalva et al., EphB receptors interact with NMDA receptors and regulate excitatory synapse formation. Cell 103, 945–956 (2000).

10. C. Minkin, Bone acid phosphatase: tartrate-resistant acid phosphatase as a marker of osteoclast function. Calcif Tissue Int 34, 285–290 (1982).

11. H. Rico, L. F. Villa, Serum tartrate-resistant acid phosphatase (TRAP) as a biochemical marker of bone remodeling. Calcif Tissue Int 52, 149–150 (1993).

12. M. J. Zylka et al., Prostatic acid phosphatase is an ectonucleotidase and suppresses pain by generating adenosine. Neuron 60, 111–122 (2008).

13. P. Guedes-Dias, E. L. F. Holzbaur, Axonal transport: Driving synaptic function. Science 366, (2019).

14. N. Ahimsadasan, V. Reddy, M. Z. Khan Suheb, A. Kumar, in StatPearls. (StatPearls Publishing Copyright © 2024, StatPearls Publishing LLC., Treasure Island (FL), 2024).

15. S. I. Sheffler-Collins, M. B. Dalva, EphBs: an integral link between synaptic function and synaptopathies. Trends Neurosci 35, 293–304 (2012).

16. A. Y. Kim et al., Pirt, a phosphoinositide-binding protein, functions as a regulatory subunit of TRPV1. Cell 133, 475–485 (2008).

17. A. Latremoliere, C. J. Woolf, Central sensitization: a generator of pain hypersensitivity by central neural plasticity. J Pain 10, 895–926 (2009).

18. S. N. Hassler, et al., The cellular basis of protease-activated receptor 2–evoked mechanical and affective pain. JCI Insight 5, (2020).

19. E. V. Brown, A. F. Malik, E. R. Moese, A. F. McElroy, A. C. Lepore, Differential Activation of Pain Circuitry Neuron Populations in a Mouse Model of Spinal Cord Injury-Induced Neuropathic Pain. J Neurosci 42, 3271–3289 (2022).

20. A. Barik et al., A spinoparabrachial circuit defined by Tacr1 expression drives pain. Elife 10, (2021).

21. S. Choi et al., Parallel ascending spinal pathways for affective touch and pain. Nature 587, 258–263 (2020).

22. A. M. Bell et al., Deep sequencing of Phox2a nuclei reveals five classes of anterolateral system neurons. bioRxiv, (2023).

23. O. Gautier et al., Challenges of profiling motor neuron transcriptomes from human spinal cord. Neuron 111, 3739–3741 (2023).

24. J. A. Blum et al., Single-cell transcriptomic analysis of the adult mouse spinal cord reveals molecular diversity of autonomic and skeletal motor neurons. Nat Neurosci 24, 572–583 (2021).

25. T. L. Daigle et al., A Suite of Transgenic Driver and Reporter Mouse Lines with Enhanced Brain-Cell-Type Targeting and Functionality. Cell 174, 465–480.e422 (2018).

26. T. Huang et al., Identifying the pathways required for coping behaviours associated with sustained pain. Nature 565, 86–90 (2019).

27. B. S. Khakh, Molecular physiology of p2x receptors and atp signalling at synapses. Nature Reviews Neuroscience 2, 165–174 (2001).

28. J. P. Dupuis, O. Nicole, L. Groc, NMDA receptor functions in health and disease: Old actor, new dimensions. Neuron 111, 2312–2328 (2023).

29. R. C. Malenka, R. A. Nicoll, NMDA-receptor-dependent synaptic plasticity: multiple forms and mechanisms. Trends Neurosci 16, 521–527 (1993).

30. B. C. Park, M. Reese, V. S. Tagliabracci, Thinking outside of the cell: Secreted protein kinases in bacteria, parasites, and mammals. IUBMB Life 71, 749–759 (2019).

31. R. Ruscheweyh, O. Wilder-Smith, R. Drdla, X.-G. Liu, J. Sandkühler, Long-Term Potentiation in Spinal Nociceptive Pathways as a Novel Target for Pain Therapy. Molecular Pain 7, 1744–8069-1747-1720 (2011).

32. C. N. Sang, NMDA-Receptor Antagonists in Neuropathic Pain: Experimental Methods to Clinical Trials. Journal of Pain and Symptom Management 19, 21–25 (2000).

33. M. Hruska, N. Henderson, S. J. Le Marchand, H. Jafri, M. B. Dalva, Synaptic nanomodules underlie the organization and plasticity of spine synapses. Nature Neuroscience 21, 671–682 (2018).

34. Y. T. Mao et al., Filopodia Conduct Target Selection in Cortical Neurons Using Differences in Signal Kinetics of a Single Kinase. Neuron 98, 767–782.e768 (2018).

35. M. R. Bordoli et al., A secreted tyrosine kinase acts in the extracellular environment. Cell 158, 1033–1044 (2014).

36. P. Chander, M. J. Kennedy, B. Winckler, J. P. Weick, Neuron-Specific Gene 2 (NSG2) Encodes an AMPA Receptor Interacting Protein That Modulates Excitatory Neurotransmission. eneuro 6, ENEURO.0292-0218.2018 (2019).

37. D. Guez-Barber et al., FACS purification of immunolabeled cell types from adult rat brain. J Neurosci Methods 203, 10–18 (2012).

38. M. Hruska, N. T. Henderson, N. L. Xia, S. J. Le Marchand, M. B. Dalva, Anchoring and synaptic stability of PSD-95 is driven by ephrin-B3. Nat Neurosci 18, 1594–1605 (2015).

39. A. J. B. Kreutzberger et al., Reconstitution of calcium-mediated exocytosis of dense-core vesicles. Science Advances 3, e1603208 (2017).

40. J. Rappsilber, M. Mann, Y. Ishihama, Protocol for micro-purification, enrichment, pre-fractionation and storage of peptides for proteomics using StageTips. Nat Protoc 2, 1896–1906 (2007).

41. Y. Xu et al., Cardiolipin remodeling enables protein crowding in the inner mitochondrial membrane. Embo j 40, e108428 (2021).

42. J. Cox et al., Andromeda: a peptide search engine integrated into the MaxQuant environment. J Proteome Res 10, 1794–1805 (2011).

43. S. Tyanova, T. Temu, J. Cox, The MaxQuant computational platform for mass spectrometry-based shotgun proteomics. Nat Protoc 11, 2301–2319 (2016).

44. R. Craig, J. P. Cortens, R. C. Beavis, Open source system for analyzing, validating, and storing protein identification data. J Proteome Res 3, 1234–1242 (2004).

45. S. D. Sherrod et al., Label-free quantitation of protein modifications by pseudo selected reaction monitoring with internal reference peptides. J Proteome Res 11, 3467–3479 (2012).

46. S. Shiers, R. M. Klein, T. J. Price, Quantitative differences in neuronal subpopulations between mouse and human dorsal root ganglia demonstrated with RNAscope in situ hybridization. PAIN 161, 2410–2424 (2020).

47. S. I. Shiers et al., Convergence of peptidergic and non-peptidergic protein markers in the human dorsal root ganglion and spinal dorsal horn. Journal of Comparative Neurology 529, 2771–2788 (2021).

48. S. R. Chaplan, F. W. Bach, J. W. Pogrel, J. M. Chung, T. L. Yaksh, Quantitative assessment of tactile allodynia in the rat paw. J Neurosci Methods 53, 55–63 (1994).

49. D. J. Langford et al., Coding of facial expressions of pain in the laboratory mouse. Nature Methods 7, 447–449 (2010).

50. M. Cheah, J. W. Fawcett, M. R. Andrews, Assessment of Thermal Pain Sensation in Rats and Mice Using the Hargreaves Test. Bio-protocol 7, e2506 (2017).

51. D. S. Brenner, J. P. Golden, R. W. t. Gereau, A novel behavioral assay for measuring cold sensation in mice. PLoS One 7, e39765 (2012).

52. A. M. Cowie, C. L. Stucky, A Mouse Model of Postoperative Pain. Bio-protocol 9, e3140 (2019).

