## Supplemental Materials and Figures for "VLK drives extracellular phosphorylation of EphB2 to govern the EphB2-NMDAR interaction and injury-induced pain"

**The PDF file includes:**

Materials and Methods
Figs. S1 to S7
Tables S1 to S2
References

**Other Supplementary Materials for this manuscript include the following:**

Movies S1

### Materials and Methods

#### Animals

The Institutional Animal Care and Use Committee of Thomas Jefferson University, Tulane University, and UT Dallas approved all animal procedures. All behavior tests were conducted during the light hours of the cycle, between 6:00 a.m. and 6:00 p.m. Animals were group-housed, with each cage having an equal balance of treatment conditions/genotypes. Dissociated cortical culture neurons were prepared from E17-18 Long Evans rat (Charles River). Wild-type CD-1 mice (Charles River) and transgenic mice (generated for and described in this study) were housed (3-5 mice per cage) in Thomas Jefferson University, Tulane University, and UT Dallas laboratory animal facilities.

#### Human Tissue

The Institutional Review Boards of UT Dallas (UTD) reviewed and approved protocols. All experiments conformed to relevant guidelines and regulations. Dorsal root ganglion tissues were collected from the donors after neurological determination of death within 2-3 hours of cross-clamp. Donor information is provided in Table S1.

#### HEK293T cell culture and transfection

HEK293T cells were cultured and transfected using calcium phosphate as described (*4*). NMDAR-transfected HEK293T media was supplemented with 50μM APV (Tocris Bioscience) and 10μM MK801 (Tocris Bioscience) to prevent excitotoxicity (*4*). Stable 293T VLK knockdown cell line was generated by transfecting cells with a shRNA targeting *PKDCC* (5'-CCCAACGTGCTGCAGCTCT-3'; Horizon Discovery Ltd.) followed by prolonged puromycin (2μg/ml) selection over 10 passages. The stable *PKDCC* knockdown clone was maintained in puromycin (0.5μg/ml) containing media.

#### Primary neuronal culture and transfection

Primary neuronal cultures were derived from embryonic day 17-18 (E17-18) rat cerebral cortex and maintained as described previously (*4, 6*). $8X10^6$ neurons were plated for 10cm dishes and $1x10^6$ neurons were plated per well of 6 well dishes for biochemical assays. $2x10^5$ neurons were plated on glass bottom dishes for live cell imaging experiments. As previously described, neuronal cultures were transfected using Lipofectamine 2000 (*4, 6*).

#### Human iPSC neuronal differentiation and maturation (hiSNs)

Human iPSC-derived immature sensory neurons (RealDRGx™) were produced by Anatomic Inc. with the Anatomic SensoX-DM kit (Anatomic #7007). Neurons at day 7 post-differentiation were plated on glass coverslips coated with poly-L-ornithine (Sigma, Castle Hill Australia) and Matrix 3 (Anatomic Inc., Minneapolis, USA) in 96-well tissue culture plates (Corning) at a density of 40,000 cells/$cm^2$, and maintained in Chrono™ Senso-MMx1 maturation medium (Anatomic cat# 7008)for 3 days followed by Chrono™ Senso-MMx2 for the remaining 28 days (DIV28). Growth media was exchanged three times a week, and stimulations were performed at DIV28.

#### Expression Constructs

Myc-GluN1, GluN2B, FLAG-EphB2, mutant FLAG-EphB2 (Y504E and Y504F), and EGFP have been previously described (*4, 6*). Syp1-EGFP and mTurquoise2 were previously described (*33, 34*). Flag-tagged constructs of VLK, VLK-Kinase dead (KD), DIA1, DIA1R, Fam69A, Fam69B, and Fam69C have been described (*1, 35*). The Flag-tagged VLK rescue construct (shRNA resistant) was created by using site-directed mutagenesis (Stratagene, La Jolla, CA) to convert the *PKDCC* shRNA target site described above to 5'- CCaAAtGTaCTaCAaCTa 3' with the following primers Fwd 5'-

CTGCTGGAGCGGCTGCGGCACCCAAATGTACTACAACTATATGGCTACTGCTACCAGGAC-3’ and Rev 5’-TCCTGGTAGCAGTAGCCATATAGTTGTAGTACATTTGGGTGCCGCAGCCGCTCCAGCAG-3’ without changing the protein sequence. The VLK-mCherry lentiviral construct was generated by PCR amplifying VLK from above and subcloning it downstream of the human synapsin promoter to replace NSG2 in pFCK-NSG2-mCherry (*36*) using standard restriction cloning.

**Lentivirus production and transduction**

Lentivirus was packaged using a second-generation packaging system by transfecting HEK293T cells with VLK-mCherry, psPAX2, and pMD2.G as previously described (*36*). Lentivirus was harvested from transfected cell media by ultracentrifugation 0.45µm filter-sterilized supernatant at 110,000xG for 2h. DIV3 neurons were transduced with VLK-mCherry lentivirus (MOI 2-3) and assayed at DIV7 for ephrinB2-dependent VLK secretion.

**Western Blotting, Immunoprecipitation (IP) and Co-IP**

HEK293T cells and cultured neurons were lysed in RIPA buffer to generate protein lysates. Cortices of postnatal day 35 (P35) transgenic mice *Pkdcc*$^{flox/flox}$ *CaMKII*$^{Cre/+}$ and its WT littermate controls were also lysed using RIPA buffer and an equal amount of protein was used to perform IP. Pooled dorsal horns from *Pkdcc*$^{flox/flox}$ *Pirt*$^{Cre/+}$ uninjured mice and their WTlittermate controls were lysed in RIPA. Following lysis, an additional Percoll gradient spin was carried out to remove myelin and unwanted cellular debris (*37*) and an equal amount of protein was used to perform IP. For conditioned ACSF IP, ACSF was filtered through a 0.22µm syringe filter (Millipore, cat#SLGV004SL) and incubated with antibody for 2h at 4$^{o}$C followed by addition of Protein-G beads and incubation overnight at 4$^{o}$C. IP, Co-IP, and Western blotting were performed as previously described (*4, 6*). RFP Trap agarose beads (ChromoTek, cat# rta) were used to IP VLK-mCherry from as per manufacturer’s instructions. For RFP Trap IP conditioned ACSF was filtered as described above and further modified by addition of a final concentration of 10mM Tris pH 7.5, NaCl 150mM, 0.5mM EDTA and 0.5% NP-40. Antibodies used for IP were mouse monoclonal Anti-FLAG M2 (Sigma, cat# F1804, lot# SLCN3722), goat polyclonal anti-EphB2 (R&D Systems, cat# AF467, lot# CVT0315041), mouse monoclonal anti-pTyr (PY99) (Santa Cruz, Cat# SC-7020; lot# L022)

**Cellular Stimulations**

Ephrin-B2 – Control or clustered ephrin-B2 treatments were performed as previously described (*9*). Briefly, control IgG-Fc (Cat#110-HG-100; R&D Systems) and recombinant ephrin-B2-Fc (Cat#7397-EB-050; R&D Systems) were clustered with Donkey anti-Human IgG (Cat#709-005-149; Jackson ImmunoResearch) for 30min at room temperature. Multimerized control and ligand were mixed in 37°C artificial cerebrospinal fluid (ACSF, 140 mM NaCl, 5 mM KCl, 1 mM MgCl2, 2 mM CaCl2, 20 mM glucose, and 10 mM HEPES, pH 7.3) and applied on cells for 45 min. The ACSF was collected from the wells to detect secreted VLK, passed through a 0.22µm filter, and immediately subjected to IP as indicated or stored at -80$^{o}$C until used.

Recombinant protein – HEK293T cells were treated with 100ng recombinant VLK, Fam69C, or DIA1. Synaptosomes were treated with 100ng recombinant VLK. 1mM ATP (Cat#A9187; Sigma) was applied as indicated. Recombinant PAP treatment was applied at 100ng.

Botulinum toxin – Active Botulinum toxin A or B (BoNTA/B)(Cat#128C and 138B (discontinued; List labs, CA) were applied at 100pM in ACSF. Cells were pretreated for 1h with toxins in ACSF followed by application of ACSF containing toxins with either control or ephrin-B2 for 45min.

**Proximity Ligation Assay (PLA)**
As previously described, PLA experiments on HEK293T cells, mouse brain, and human spinal cord synaptosomes were performed (*4*). Synaptosome PLA combined with vGlut1 immunostaining was also previously described (*4*).

***In vitro* Kinase assay**
For the *in vitro* kinase assay purified recombinant protein fragment with EphB2 extracellular domain (EphB2 (200ng), active; Millipore) and active VLK (100 ng), VLK-KD (100ng), Fam69C (100ng), DIA1(100ng) was used. The kinase reaction was started by adding 10 mM MnCl2, 10 mM MgAc, and 100 µM ATP in ACSF buffer. After 30 min of incubation at 37 °C, phosphorylated proteins were separated by SDS-PAGE and analyzed by western blotting with anti-phosphotyrosine antibody (PY99) and EphB2. This protocol was also previously described (*6*)

**RNA extraction and qPCR**
Total RNA from HEK293T cells was extracted using TRI reagent (Cat#TR118; Molecular Research Center Inc.) as per the manufacturer's instruction. Total RNA from mice cortices was extracted using the Direct-zol RNA microprep kit (Cat#R2062; Zymo Research) following the manufacturer's instructions. RNA was quantified using a Nanodrop and 1ug of total RNA was used to prepare cDNA using qScript cDNA Synthesis kit (Cat# 90547-100; Quantabio) as per manufacturer's instructions for quantitative polymerase chain reaction (qPCR). PCR was set in triplicates using SYBR green (Applied Biosystems) in a 20µL volume and using 15 ng cDNA template. PCR was run using the StepOne Plus system, and the quantification was performed using the comparative Ct (ΔΔCt) method. The qPCR primers used are described in Table S2.

**VLK, Fam69C, DIA1, and VLK-KD Protein Purification**
GST-tagged kinase-containing plasmids were transformed into competent BL21 cells. A single colony was picked and grown overnight in 5 mL LB-ampicillin starter culture. 1 Liter LB-ampicillin cultures were inoculated and grown at 37°C and 250 rpm until OD600 reached 0.8-1. Cultures were cooled to 16°C. 0.03 mM IPTG (optimized for VLK) was added. The cultures were induced overnight at 16°C. A 2 mL glutathione resin slurry was loaded into a column and washed with 5 mL resuspension buffer by gravity flow. The column was stored in a resuspension buffer at 4°C for next-day use. The following day, the bacteria were pelleted at 6000 rpm for 10 minutes, and the pellet was resuspended in 35 mL resuspension buffer. Lysozyme and PMSF were added and mixed. Lysate was incubated on ice for 10 min. Lysate was sonicated 20 X 3 sec ON, 3 sec OFF on ice using a probe sonicator. Lysate was centrifuged at 15,000 rpm (~28,000 X g) for 20 min at 4°C. The supernatant was transferred to a clean container immediately. The supernatant was loaded slowly onto the glutathione column. The column was washed 4X with wash buffer, 5 mL each. The column was plugged, and 4mL elution buffer was added. Beads were suspended and incubated for 10 min. The 4 mL elution was collected, and another 2 mL elution buffer was added to the column and collected. Protein elution was then dialyzed overnight.

**Brain Fractionation**
<u>Synaptosomes</u> – Synaptosomes from wild type CD-1 mice brains were prepared and plated as previously described (*4, 6*).

<u>Post-synaptic Density (PSD) fractionation</u> - PSD fractions were prepared from postnatal day 21 (P21) wild-type CD-1 mice (*38*). Brains were homogenized on ice in 0.32 M sucrose, 4 mM HEPES, pH 7.4, containing a protease inhibitor cocktail (Sigma) and 1 mM PMSF. After removing the nuclear fraction (P1) by centrifugation at 1,000 x g for 15 minutes at 4°C, non-synaptic fractions were further centrifuged at 10,000 x g at 4°C to obtain the crude synaptosomal fraction (P2). This pellet was resuspended in 10 volumes of HEPES-buffered sucrose and then spun again

at 10,000 x g for another 15 min. The resulting pellet was lysed by hypo-osmotic shock in water, rapidly adjusted to 4mM HEPES pH 7.4, and mixed constantly for 30 min at 4°C. The lysate was centrifuged at 25,000 x g for 20 min and the pellet (P3) was resuspended in HEPES-buffered sucrose. The crude synaptic vesicle supernatant (S3) was saved for further purification of synaptic vesicles. The resuspended membranes were carefully layered on a discontinuous gradient containing 0.8 to 1.0 to 1.2 M sucrose and centrifuged at 150,000 x g for 2h. Synaptic plasma membranes were recovered in the layer between 1.0 and 1.2 M sucrose and diluted to 0.32 M sucrose by adding 2.5 volumes of 4 mM HEPES pH 7.4. Membranes were pelleted by centrifugation at 150,000 x g (36,000 rpm in SW50.1) for 30 minutes. Pellet (SPM) was resuspended in 3-5 ml of ice-cold 50 mM HEPES pH7.4, 2mM EDTA, plus protease/phosphatase inhibitors. Triton X-100 was added to 0.5%. Samples were rotated in the cold room for 15 min. Samples were centrifugated at 32,000 × g (16,318 rpm in SW50.1) for 20 min to obtain the PSD-1T pellet. Pellet was resuspended in 3 mL ice-cold 50 mM HEPES pH7.4, 2 mM EDTA plus protease/phosphatase inhibitors). 1 mL was saved as PSD-1T. To half of the remaining resuspended pellet (~l mL), Triton X-100 was added to 0.5% and rotated in the cold room for 15 minutes. Sample was centrifugated at 200,000 × g (41,000 rpm in SW50.1) for 20 min to obtain the PSD-2T pellet. In a separate experiment, resuspend the second half of the PSD-IT pellet and incubate for 10 min in ice-cold 3% sarcosyl (N-lauroyl sarcosine) in 50 mM HEPES pH7.4, 2 mM EDTA, plus protease/phosphatase inhibitors. Centrifuge at 200,000 × g (41,000 rpm in SW50.1) for 1 hour to obtain the PSD-S pellet. Pellets were resuspended in 50 mM HEPES pH7.4, 2 mM EDTA plus protease/phosphatase inhibitors.

Dense Core Vesicles Isolation (DCV) - DCVs were prepared from postnatal day 21 (P21) wild-type CD-1 mice (*39*). Brains were homogenized on ice in a homogenization medium (0.26 M sucrose, 5 mM MOPS, and 0.2 mM EDTA). Homogenate was then spun at 4000 rpm (1000 × g) for 10 min at 4°C in a fixed-angle microcentrifuge to pellet nuclei and larger debris. The post nuclear supernatant (PNS) was collected and spun at 11,000 rpm (8000 × g) for 15 min at 4°C to pellet mitochondria. The post-mitochondrial supernatant (PMS) was then collected, adjusted to 5 mM EDTA, and incubated for 10 min on ice. A working solution of 50% OptiPrep (iodixanol) [five volumes of 60% OptiPrep (one volume of 0.26 M sucrose, 30 mM MOPS, and 1 mM EDTA)] and homogenization medium were mixed to prepare solutions for discontinuous gradients in Beckman SW 55 tubes: 0.5 ml of 30% iodixanol on the bottom and 3.8 ml of 14.5% iodixanol, above which 1.2 ml of EDTA-adjusted PMS was layered. Samples were spun at 45,000 rpm (190,000 × g) for 5 hours. 10 fractions were collected and analyzed by Western blot. A clear white band at the interface between the 30% iodixanol and the 14.5% iodixanol was the DCV sample (fraction 9 of Western blot).

**Proteomic analysis**
Proteins samples from S3 crude synaptic vesicle fraction from 3 male and 3 female mice were concentrated by short SDS-PAGE runs into single-protein bands before tryptic digestion and peptide extraction. Peptides were desalted with Empore C18 High Performance Extraction Disks (*40*), and the eluted peptide solutions were partially dried under vacuum and then analyzed by LC-MS/MS with a Thermo Easy nLC 1000 system coupled online to a Q Exactive HF with a NanoFlex source (Thermo Fisher Scientific) as previously described (*41*). All data were analyzed by MaxQuant proteomics software (version 1.5.5.1) with the Andromeda search engine (*42, 43*) using a mouse protein FASTA (mouse [Mus musculus] protein database; Uniprot; Reviewed, 21,989 entries [06182020]). Additionally, targeted mass spectrometry analysis was carried out to identify extracellular tyrosine-protein kinase (VLK) in both female and male preparations. Theoretical monoisotopic masses of 6 tryptic peptides of VLK that are commonly seen and documented in the Global Proteome Machine resource (GPM, (*44*)) were used in an inclusion list. The resulting raw mass spectrometry files were imported into Skyline proteomics software (*45*),

and the presence of precursor and transition ions confirmed to identify VLK in both male and female peptide pools unequivocally.

**Live cell imaging, Immunocytochemistry and Image analysis**

Live imaging was carried out on rat cortical neurons plated on 35mm glass bottom dishes (Cell E&G). DIV17 cultured rat cortical neurons were co-transfected with VLK-mCherry, Syp1-EGFP, and mTurquoise2. Live cell imaging was carried out on DIV21-22 neurons by replacing cell media with 37ºC ACSF for the duration of live imaging. Time-lapse images of live neurons were acquired on the Leica SP8 using a 63x objective lens. Images of neuronal axons (determined by the mTurquoise cell fill) were acquired at 2 frames/sec for 5 min as a single focal plane. Laser power was kept at low levels to avoid photobleaching. After live imaging, cells were immediately fixed with 4%PFA/2% sucrose for 8 minutes for posthoc immunocytochemistry as previously described (*4*). PSD95 primary antibody was applied overnight at 4ºC. PSD95 signal was detected with an AF647 conjugated secondary antibody. Permanent markers of different colors were used on the glass dish for reference of reimaging. Additionally, the location of each imaged axon was recorded using a computer-controlled stage. These recorded references were used to re-locate the same axons reimaged to visualize PSD95, VLK-mCherry, and Syp1-EGFP (Posthoc). Posthoc fixed cell confocal images were acquired as Z stacks at 0.4-0.6 mm intervals.

Images were analyzed using Fiji (ImageJ). A segmented line was drawn along the axon segments to measure axon length and generate kymographs using the reslice tool. On the kymographs, puncta were categorized into 2 groups – Stationary and Moving. Moving puncta were distinguished as tilted straight lines. Stationary puncta were vertical straight lines that did not deviate from the first to the last frame of live imaging. Stationary, moving, and colocalized puncta were manually scored. The number of puncta in each category was divided by the total length of the axon to derive puncta density. For posthoc analysis, images were background subtracted, gaussian blurred, and registered with live images using Turboreg (*33, 34*). The straightened axon segments from posthoc image were added as a new frame to the live image after the last frame of the identical axon segment from live imaging to assess and determine which stationary puncta were maintained after fixation. The stationary VLK-mCherry and Syp1-EGFP puncta maintained after fixation were manually scored. A custom macro (*4, 34*) was used to determine colocalization of maintained puncta with each other and PSD95. The analysis did not include Axon segments with no stationary puncta after fixation. Moving VLK puncta velocities were determined by manually tracking moving puncta using the MTrackJ plugin.

**DRG dissection, RNAScope, Immunohistochemistry**

DRG from Mouse and Human donors were processed as previously described (*46*). Immunohistochemistry and RNAScope were carried out on mouse and human DRG as previously described (*46*). Briefly, mRNA transcripts were detected using the RNAscope Fluorescent Multiplex Assay (Advanced Cell Diagnostics) and RNAscope Fluorescent Multiplex Reagent Kit (#320851 & #323100). The RNAscope catalogue probes used to detect were (For mouse *Pkdcc* (#516961), *Scn10a* (#426011-C2), *Tacr1* (#428781-C3) and human *PKDCC* (#525121), *SCN10A* (#406291-C2)). Counterstaining for NF200 and NeuN was done with immunohistochemistry on the same tissues. Tissues were blocked in 0.1M Phosphate Buffer with 10% Normal Goat Serum and 0.3% Triton-X. Primary antibody was incubated overnight at 4ºC. Secondary antibodies were incubated for 1 hour at room temperature. Coverslips were mounted with Prolong Gold Antifade (Fisher #P36930). Images were taken on an Olympus Fluoview FV1000 confocal microscope at 20X magnification. Histological analyses were carried out using Olympus Cellsens software. Frequency distribution, colocalization data and diameter measurements were calculated as described before (*47*). The Cellsens Count and Measure tool was used to conduct colocalization and cell area analyses in the DRG and spinal cord. To determine the expression-based distribution of *PKDCC* transcripts in the DRG neurons, expression levels were defined as Low (1 – 10 puncta) and High (10+ puncta).

cFos and NeuN staining was conducted on free-floating tissue sections. Adult mice were euthanized by isoflurane inhalation. Lumbar spinal cords were dissected and drop fixed in 10% formalin for 2 hours, cryoprotected, embedded in OCT (Fisher #1437365) and frozen on dry ice. Tissues were cryostat sectioned at 40 μm. Tissues were blocked in 0.1M Phosphate buffer with 10% Normal Goat Serum and 0.3% Triton-X and incubated in cFos and NeuN primary antibodies for 48 hours at 4°C. Secondary antibodies were incubated for 2 hours at room temperature. Sections were slide mounted and cover slipped with Prolong Gold Antifade. Images were taken on an Olympus Fluoview FV1000 confocal microscope at 20X magnification. $cFos^+$ cell counts were carried out using Imaris software.

**Generation of a VLK-specific Antibody**

Antibodies against the ATP binding region of VLK were generated in rabbits against a peptide of the sequence Ac-CKALKAVDFSGHDLGS-$NH_2$ (Covance, Denver, PA). Serum was collected and passed through a column containing SulfoLink Coupling Resin (Thermo Scientific) conjugated to KLH peptide. The serum was then incubated overnight in a column containing resin conjugated to the above VLK peptide. The antibody was eluted with glycine pH 2.0 and then extensively dialyzed in PBS.

**Somascan Assay**

Tissue lysates were prepared from fresh-frozen DRG, and from the dorsal and ventral horn portions of the lumbar spinal cord. The tissues were placed in T-PER Tissue Protein Extraction Reagent (Thermo Scientific, Cat # 78510) with additional 1X Halt Protease Inhibitor Cocktail (Thermo Scientific, Cat # 87786) and homogenized using Precellys Soft Tissue Homogenizing beads (Bertin Corp, Cat # P000933-LYSK0-A.0). Samples were centrifuged at 14,000 *x g* for 15 minutes in the cold room. The resulting supernatant was quantified Micro BCA™ Protein Assay Kit (Thermo Scientific, Cat# 23235) and normalized accordingly. Proteins were profiled using the SOMAScan platform. 7000 analytes were measured on the SOMAScan assay. The quality controls were performed by SomaLogic to correct technical variability within and between runs for each sample.

**Generation of *Pkdcc*$^{foxl/flox}$ mice and respective cKO mice lines**

*Pkdcc*-floxed mice were generated at the UT Health San Antonio Mouse Genome Engineering Facility via CRISPR/Cas9-mediated HDR in zygote. Briefly, two DSBs flanking the first exon and upstream enhancer sites of the *Pkdcc* gene were produced and LoxP casettes were introduced via homologous recombination (shown in Fig. 2).
The recombination induced by Cre-recombination of exon 1 would result in causing *Pkdcc* loss of function. Thus, resulted mice were bred, and homozygous *Pkdcc*$^{flox/flox}$ was selected by PCR genotyping. For genotyping, DNA was extracted from the ear samples during weaning to determine the genotypes of *Pkdcc* flox mice (KAPA mouse genotyping kit). The primers used for genotyping were mentioned in Table S2.The PCR products were visualized by electrophoresis on a 2-3% agarose gel.The *Pkdcc* floxed mice were bred with *CaMKII-Cre* mice to delete *Pkdcc* in excitatory neurons. The *Pirt*-Cre transgenic mouse line in this study was obtained from Xinzhong Dong at Johns Hopkins University (C13783-*Pirt*-Cre mice) and used to generate sensory neuron-specific knockout of *Pkdcc*. To determine if VLK is acting on EphB2 receptors expressed on these neurons, we generated a mouse line expressing LoxP sites flanking the third exon of the *Ephb2*

gene. These mice were crossed with the *Tacr1*-Cre driver line to create conditional knockouts of EphB2 in spinal cord projection neurons.

**Behavior**

Mechanical Withdrawal Thresholds were determined using von Frey Filament testing, following the method of (*48*). Animals were habituated to plexiglass chambers (11.4 × 7.6 × 7.6 cm) suspended on wire mesh racks. Baseline withdrawal thresholds were measured before experimental treatments, and subsequent testing was done at various timepoints indicated in each figure. Mechanical sensitivity was determined by applying von Frey filaments to the plantar aspect of the hind paws, and a response was indicated by flicking or withdrawal of the paw. For mice subjected to hindpaw incision surgeries, filaments were instead applied to the heel of the hindpaw near the site of injury but avoiding scar tissue.

Mouse grimace scoring was conducted using the Mouse Grimace Scale (MGS) described by (*49*). After acclimating to the suspended plexiglass chambers for an hour, mice were scored according to the MGS by blinded experimenters at baseline and post treatment time points which are indicated in each figure. The grimace scores were averaged by group at each time point and plotted respectively.

Intrathecal injections were performed as previously described under isoflurane anesthesia (*6*). Drugs were administered in a 5 uL volume with a Hamilton syringe. Drugs were diluted in sterile saline. Heat denaturation was done by heating proteins at 70°C for 30 mins. VLK, KD-VLK, DIA1, FAM69c and denatured controls were administered at concentrations of 0.1mg/mL PAP was administered at 250mU (*12*). APV was administered at 0.2mg/mL.

Radiant heat sensitivity was determined using the Hargreaves method (*50*). Mice were placed on a warmed glass floor (29˚ C) 20 minutes before each testing and, using a Hargreaves apparatus (IITC Model 390), a focused beam of high-intensity light was aimed at the plantar nonglabrous surface of the hind paws. The intensity of the light was set to 30% of maximum with a cutoff value of 20 seconds. The latency to withdraw either hind paw was measured to the nearest 0.01 seconds. The withdrawal latencies for both paws were averaged for each animal.

Heat tolerance/sensitivity was measured using the Hot Plate assay. A day prior to testing, mice were habituated to the Hot Plate Plexiglass box (IITC) for 10 minutes. For testing, mice were placed individually on the hot plate, heated to either 50ºC or 55ºC, and the latency to first sign of paw licking or jumping was recorded. The experiment was iterated 3 times with at least 48 hours between measurements. Stimulus cutoffs of 45 sec and 1 min were used respectively for 50ºC and 55ºC.

Cold sensitivity was measured using the cold plantar assay (*51*). Mice were habituated for 30 min to plexiglass chambers with breathing holes resting atop a 1/8th in thick glass base. The cold probe was made of finely crushed dry ice packed into a modified 3mL BD syringe. Mice were tested by pressing the cold probe against the glass directly underneath the center of the hindpaw, and latency to withdraw the paw from the cold glass was recorded with a maximum cutoff time of 20 sec. Mice were tested 4 times with at least 15 min between trials.

Sensorimotor evaluation was done using the rotarod test. Mice were acclimated to the Rotarod apparatus (IITC) a day before testing for 5 min at 4 RPM. For testing, mice were placed on the rotating rod and the RPM increased from 4 to 40 at a constant rate across 5 min. Latency to fall, RPM at fall and total distance traveled were all recorded automatically by the apparatus. 3 trials were conducted, with at least 15 minutes between trials.

Hind-paw incision surgeries were performed on transgenic animals as previously described (*6, 52*). Briefly, a ~5mm incision was made on the plantar skin of the left hindpaw under isoflurane anesthesia. The flexor digitorum brevis muscle was briefly lifted (not cut) with forceps, and the incision site was sutured closed.

Sensorimotor evaluation was done using the rotarod test. Mice were acclimated to the Rotarod apparatus (IITC) a day before testing for 5 min at 4 RPM. For testing, mice were placed on the rotating rod, and the RPM increased from 4 to 40 at a constant rate across 5 min. Latency to fall, RPM at fall, and total distance traveled were all recorded automatically by the apparatus. 3 trials were conducted, with at least 15 min between trials.

**Antibodies**

The following primary antibodies and dilutions were used:

mouse monoclonal (IgG1) anti-GluN1 (1:500 (WB), 1:200 (PLA), BioLegend, clone R1JHL, cat# 828201, lot# B383911), goat polyclonal anti-EphB2 (1:1200 (PLA), R&D Systems, cat# AF467, lot# CVT0315041), mouse monoclonal (IgG1) anti-EphB2 (1:500 (WB), Thermo Fisher Scientific, clone 1A6C9, cat# 37-1700, lot# RD2115698), mouse monoclonal (IgG2b) anti-pTyr (PY99) (1:2000 (WB) ,Santa Cruz, Cat# sc-7020; lot# L022), rabbit polyclonal anti-FLAG (1:1000 (WB), Sigma, Cat #F7425, lot# 078M4886V), mouse monoclonal (IgG2a) anti-PSD-95 (1:2500 (WB), 1:250(ICC), Neuromab, clone 28/43, cat# 75-028, lot# 472-2JU/02), mouse monoclonal (IgG1) anti-NeuN (2μg/mL (IHC), Millipore, clone A60, cat# MAB377), mouse monoclonal (IgG1) anti-Neurofilament 200 (2μg/mL (IHC), Millipore, clone N52, cat# MAB5266), guinea pig recombinant monoclonal (IgG2κ) anti-c-Fos (1:10000 (IHC), Synaptic Systems, cat# 226308), mouse monoclonal (IgG1) anti-Synaptophysin-1 (1:5000 (WB), Synaptic Systems, clone 7.2, cat# 101 111, lot# 1-46), mouse monoclonal (IgG1) anti-Synaptotagmin-1 (1:2000 (WB), Synaptic Systems, Cat# 105011, lot# 1-47), rabbit polyclonal anti-Synaptotagmin-5/9 (1:2000 (WB), Synaptic Systems, Cat# 105053, lot# 1-8), rabbit polyclonal anti-RFP (1:2000(WB), Rockland, Cat# 600-401-379, lot# 46510), mouse monoclonal (IgG1) anti-GAPDH (1:500 (WB), Millipore, Cat# MAB374, lot# 4045089), rabbit polyclonal anti-Tubulin (1:10,000 (WB), Abcam, Cat# ab18251, lot# GR235480-2), rabbit polyclonal anti-Flotilin (1:1000, Novus Bio, Cat# NBP187498, lot# 000020065), mouse monoclonal (IgG2b) anti-actin (1:2000 (ICC), Biolegend, clone 2F-1, Cat# 643801, lot# B266258), anti-Rabbit VLK (1:2000 (WB), Generated by our lab), anti-rabbit EphB2-pY662 (1:2000 (WB), previously described (*9*)), guinea pig polyclonal anti-vesicular glutamate transporter 1 (vGlut1; 1:5000 (ICC), Millipore, Cat# AB5905, lot# 3987239).

The following secondary antibodies were used:

Donkey anti-mouse-HRP (1:10,000 (WB), Jackson ImmunoResearch, cat# 715035151 lot# 151303), Donkey anti-rabbit-HRP (1:10,000 (WB), Jackson ImmunoResearch, cat# 711035152, lot# 163128), Goat anti-guinea pig DyLight-488 (1:5000 (PLA), Abcam, cat# ab102374, lot# 1041504), Donkey anti-mouse AlexaFluor-647 (1:500 (ICC), Jackson ImmunoResearch, cat# 715605150, lot#160478), Goat anti- Mouse IgG1 AlexaFluor-555 (1:500 (IHC), Thermo Fisher Scientific, cat# 21127), Goat anti-Mouse IgG1 AlexaFluor-647 (1:500 (IHC), Thermo Fisher Scientific, cat# 21240), Goat anti-Guinea pig AlexaFluor-647 (1:500 (IHC), Thermo Fisher Scientific, cat# A21450).

**Statistical analysis**

GraphPad Prism Software was used to carry out statistical analyses and generate graphs. The respective figure legends indicate the number of subjects used and the statistical analysis, including posthoc comparison tests, used for individual experiments. Mice were randomly allocated to experimental groups when possible. Behavioral testing was carried out by investigators blinded to treatment or genotype, and different experimenters successfully reproduced the results. P values of less than 5% were considered statistically significant, with * representing $P < 0.05$, ** representing $P < 0.01$, *** representing $P < 0.005$, and **** representing $P < 0.001$ unless otherwise indicated. All the models were made using Biorender.

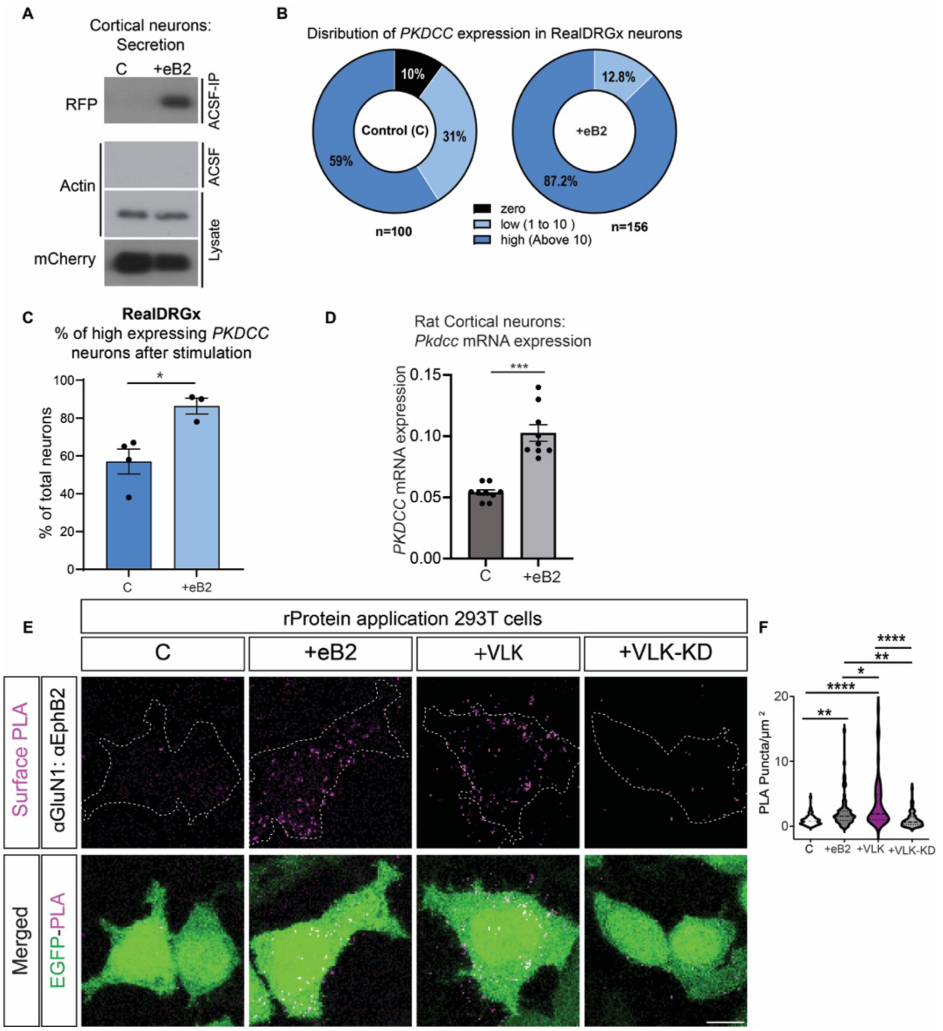


**Fig. S1. Ephrin stimulates VLK secretion, and the kinase activity of VLK is necessary to induce the EphB2-NMDAR interaction. (A)** VLK mCherry lentivirus-infected cultured cortical neurons (DIV7-8) were treated with clustered EphrinB2 (+eB2) or control (C) reagents in ACSF for 45 min. RFP Trap agarose beads was used to immunoprecipitate conditioned ACSF and immunoblotted with a custom-generated RFP antibody (ACSF-IP, top). Conditioned ACSF (ACSF, middle) and cell lysates (Lysate) were probed with beta-actin as a control to show that the conditioned media was free of any intracellular protein. mCherry was probed as lysate control (Lysate, Bottom blot). **(B)** *PKDCC* expression in Control (C) and Ephrin-treated Real DRGx neurons. No expression is depicted as Zero (black), Low is 1 to 10 puncta of *PKDCC,* and High is above 10. **(C)** Percentage of *PKDCC* expressing neurons after Control or ephrin stimulation of real DRGx neurons. **(D)** *Pkdcc* mRNA expression in rat cortical neurons.(Unpaired t-test, ***pvalue= 0.001). **(E)** Representative images of PLA of EphB2-NMDAR after treatment with soluble recombinant kinases. HEK293T cells were transfected with GluN1, GluN2B, EphB2, and EGFP. Cells were treated with Control, ephrin-B2, active recombinant VLK, and VLK-KD proteins (as indicated) for 45 minutes before fixation. The upper panels show the surface PLA signal alone (magenta). The lower panels are the merged images with EGFP in green. Scale bar = 10 µm. **(F)** Quantification of the PLA puncta number. PLA puncta numbers are quantified by counting the puncta per 100 µm$^2$ in EGFP$^+$ cells and normalized to the Control. (One way ANOVA followed by Tukey's test, **p=0.0050, 0.008, *p=0.01, ****p<0.0001, n = 70 fields from 7 repeats).

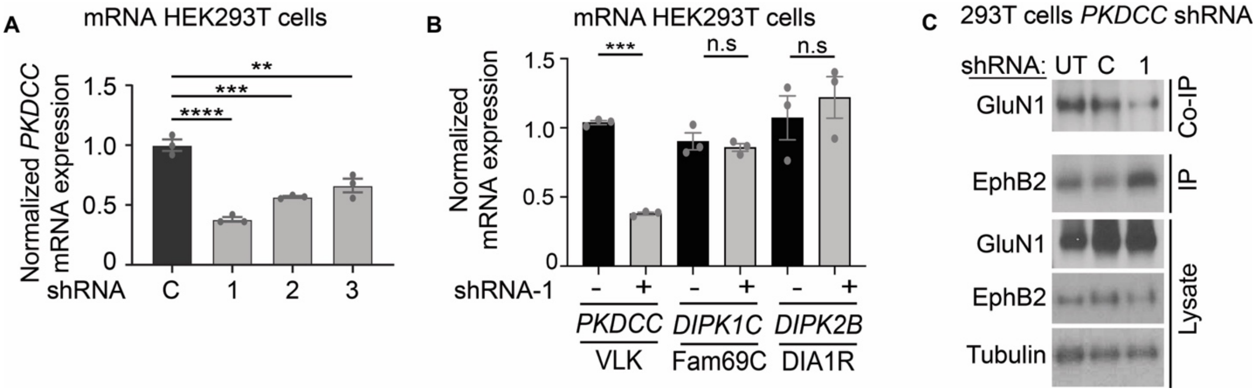


**Fig. S2. VLK knockdown abolishes EphB2-NMDAR interaction in HEK 293T cells.**
**(A)** *PKDCC* mRNA expression in HEK 293T cells following treatment with Control(C) or *PKDCC* targeting shRNA 1,2 and 3 respectively, (One way ANOVA followed by Tukey's test , ****p<0.0001, ***p=0.0002,**p=0.0012, n=3) **(B)** Real-Time (RT) PCR-based quantification of relative mRNA levels of *PKDCC* (VLK), *DIPK1C* (Fam69C), and *DIPK2B* (Dia1R) from the Stable VLK KD cells (1) (-/-, grey bars) compared to control cells (+/+, black bars). (***p<0.001, unpaired t-test; n=3). **(C)** Immunoblotting for co-immunoprecipitated GluN1 with EphB2 from HEK293T cells transfected with Untreated (UT), Control (C), and VLK shRNA1 (1) (top blot). A fraction of the same EphB2 immunoprecipitated sample was immunoblotted for EphB2 (EphB2, IP). HEK293T cell lysates were probed as indicated for EphB2 and GluN1, and Tubulin (Lysate).

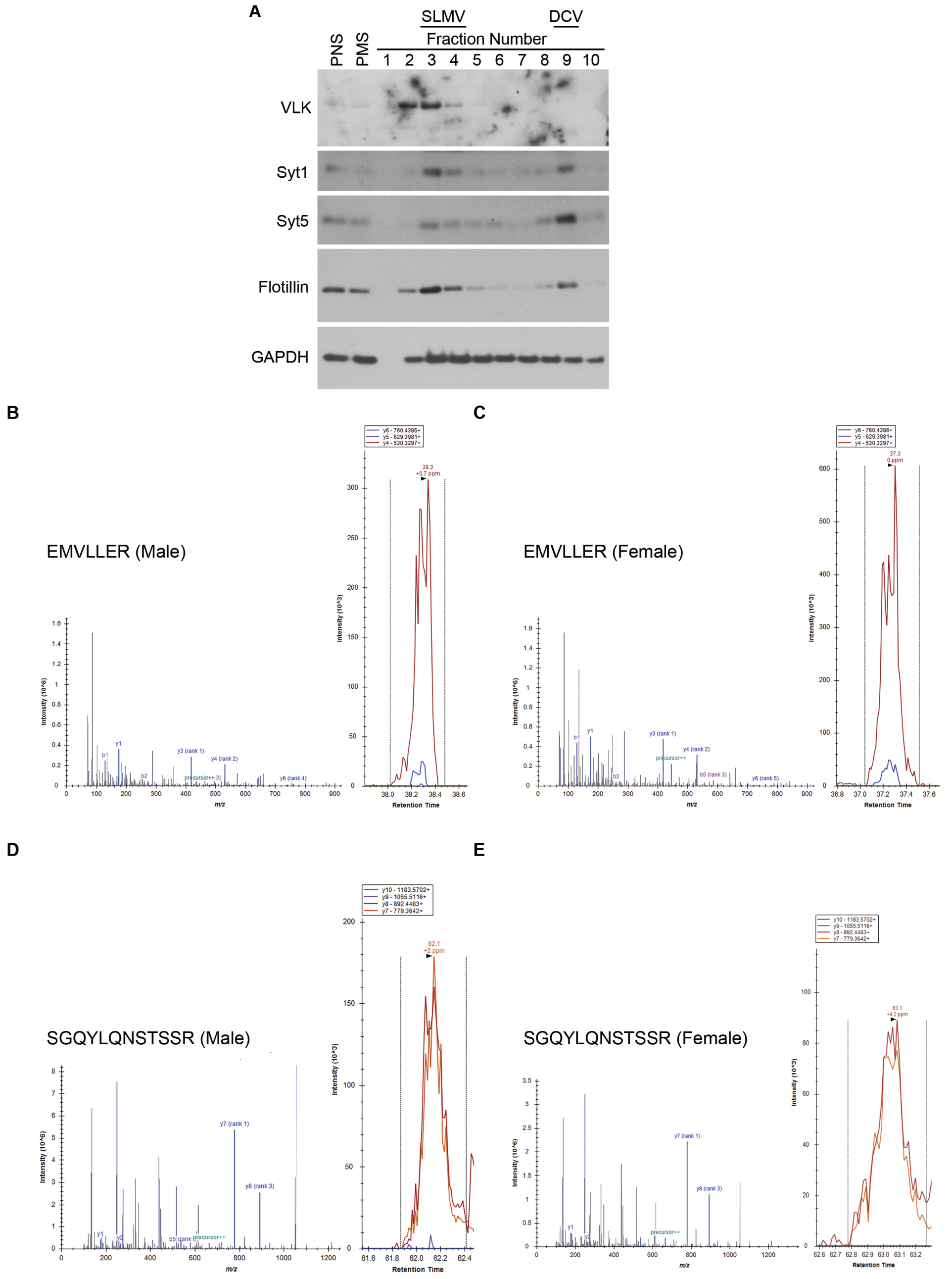


**Fig. S3. VLK is enriched in the Synaptic vesicle fraction and not in the Dense Core Vesicles. (A)** Western blots of Dense Core Vesicle purification fractions prepared from WT CD1 mouse brains showing VLK is not enriched in DCVs,but is enriched in Synaptic-like micro vesicles (SLMV). Post-nuclear supernatant (PNS), Post-mitochondrial supernatant (PMS), and Sucrose gradient fractions (1-10). Blots probed (from top to bottom): VLK, Synaptotagmin1, and Synaptotagmin 5 as DCV marker, flotillin as exosome marker, and GAPDH as a loading control. **(B-E)** Parallel Reaction Monitoring (PRM) experiment confirming the presence of VLK *(PKDCC)* and corresponding tandem MS/MS spectra for 2 of the peptides isolated and monitored are shown. Extracted ion chromatograms for major MS/MS fragment ions from the two tryptic peptides studied are shown for **(B-C)** peptide EMVLLER at *m/z*: 445.2442($2^+$) male and female, respectively, and **(D-E)** peptide SGQYLQNSTSSR at *m/z*: 664.3155 ($2^+$) male and female, respectively.

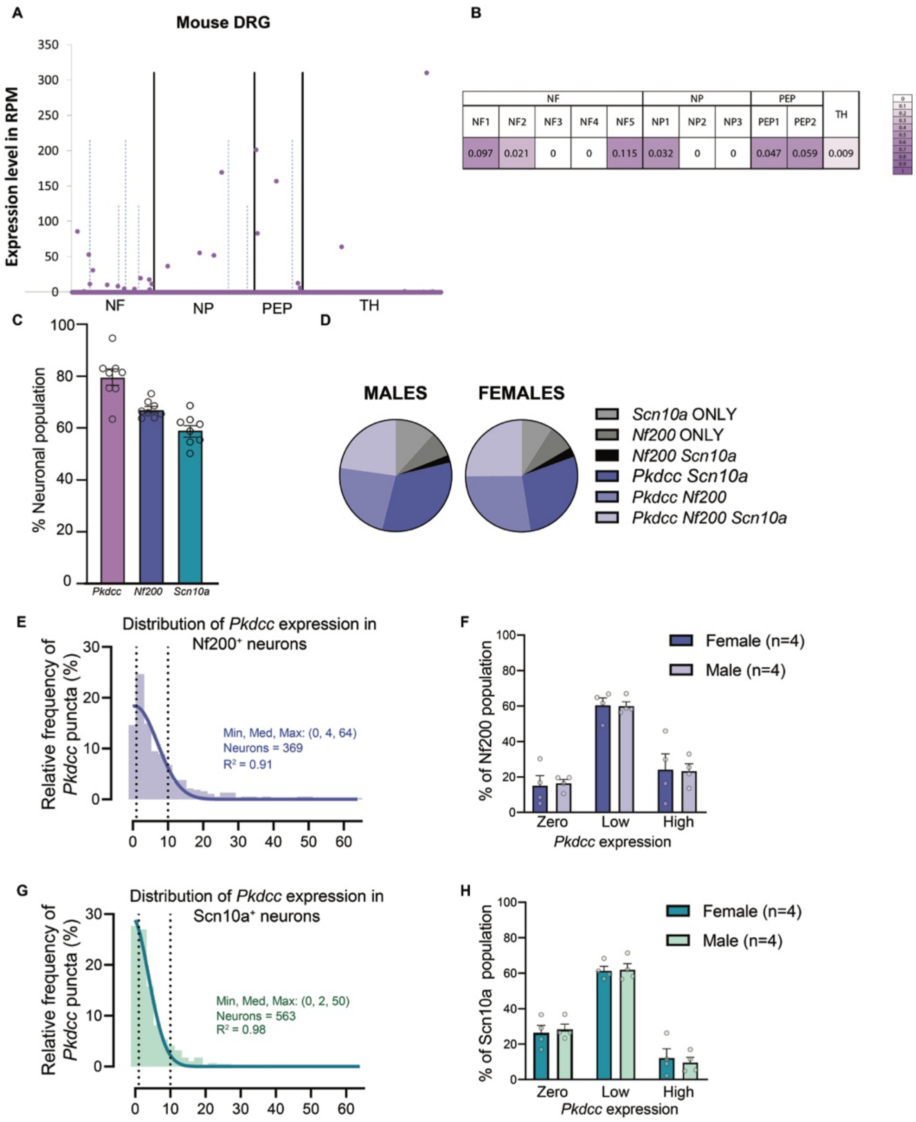

**Fig. S4.VLK expression in mouse DRG and pain circuit.**
**(A)** Cell-type expression of *Pkdcc* in the sensory neurons of mouse DRG. Solid vertical lines separate major cell populations, and dashed lines show the separation into further subtypes. Data from Linnarssonlab.org/drg/ (**B**) Quantified representation of (A) delineating the fraction of each subtype displaying a positive expression of *Pkdcc*. Data from Linnarssonlab.org/drg/. **(C)** *Pkdcc* mRNA was detected in a significant number of DRG neurons (n=8, *Pkdcc* M=79.64, SEM = 3.154; *Nf200* M = 67.15, SEM = 1.156, *Scn10a* M = 59.31, SEM = 2.048). **(D)** Population distributions were similar between male and female mice. **(E)** Histogram showing the distribution and descriptive statistics of *PKDCC* expression in Nf200$^+$ neurons. The area between the two dotted lines (1 and 10 puncta) indicates low *Pkdcc*-expressing neurons. **(F)** Sex-based analysis of *Pkdcc* expression in the Nf200 population shows no difference between males and females. **(G)** Histogram showing the distribution and descriptive statistics of *Pkdcc* expression in Scn10a$^+$ neurons. The area between the two dotted lines (1 and 10 puncta) indicates Low PKDCC-expressing neurons. **(H)** Sex-based analysis of *Pkdcc* expression in the *SCN10A* population shows no difference between males and females.

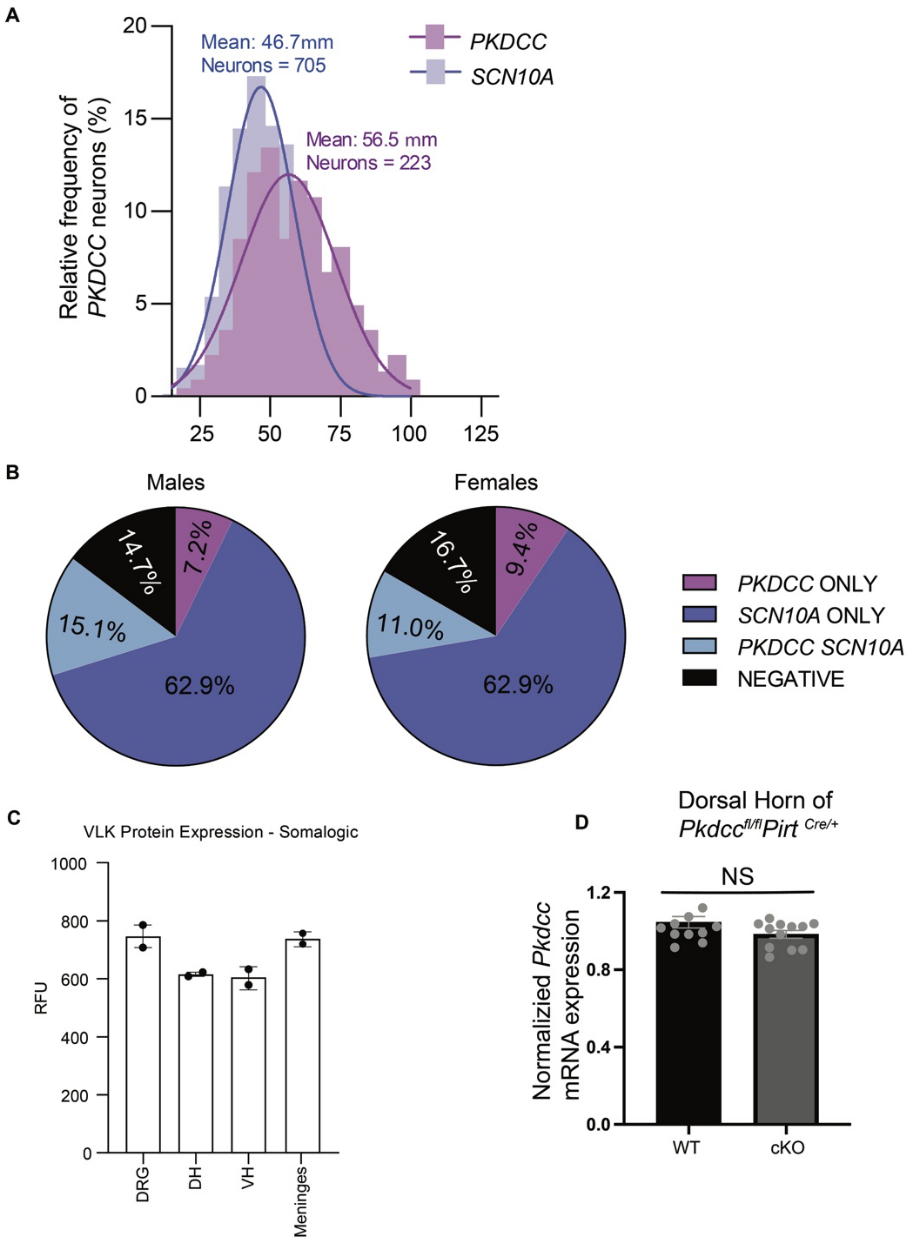


**Fig. S5. VLK expression in human DRG.**
**(A)** The Size-distribution curve of PKDCC neurons shows greater expression in small to medium-diameter cells. (**B**) *PKDCC* expression and distribution across marker populations in the human DRG is similar across males and females. **(C)** Relative Fluorescence Unit (RFU) of bulk VLK expression in different human sensory tissues using the Somalogic system, showing significant detection of VLK across the DRG, Dorsal Horn (DH), Ventral Horn (VH), and spinal meninges. **(D)** *Pkdcc* mRNA expression in WT and cKO (*Pirt*-Cre) mice.

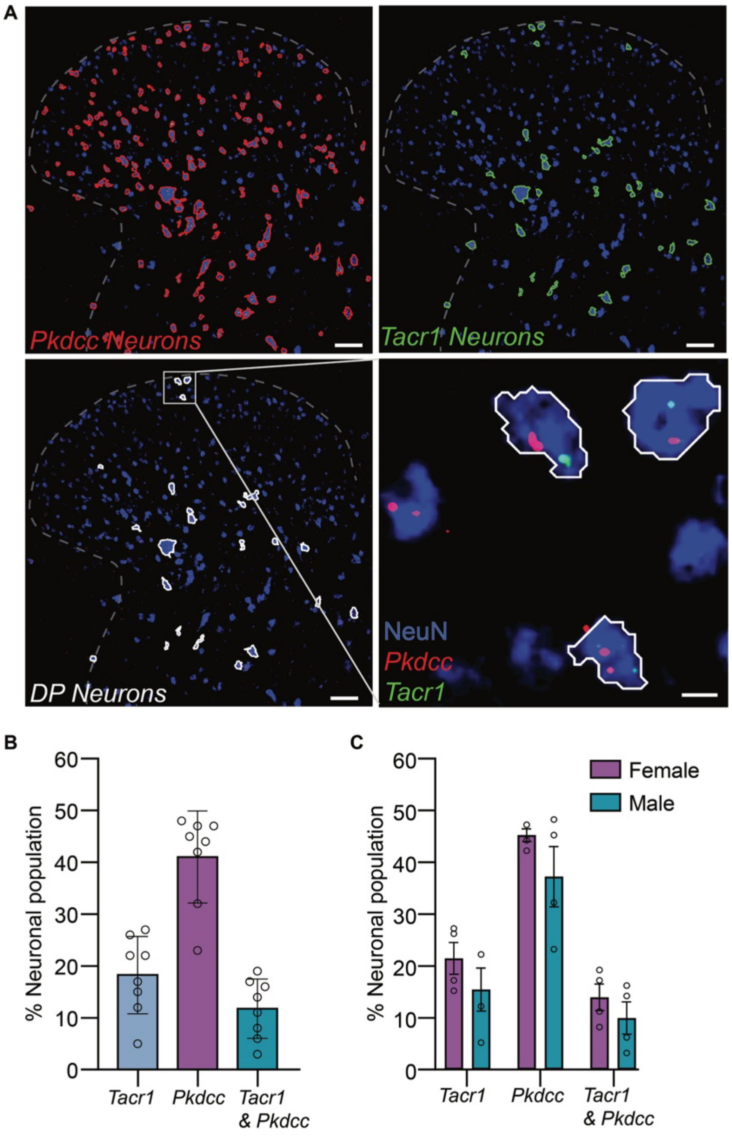


**Fig. S6. *Pkdcc is* expressed in mouse lumbar spinal cord dorsal horn neurons.**
**(A)** RNAscope *in situ* hybridization with NeuN counterstaining was used to identify *Pkdcc* and Tacr1-expressing neurons. Scale bar = 50 μm. The inset shows double-positive neurons indicated by the white borders. Scale bar = 5 μm. **(B)** Percent of DH neurons expressing *Pkdcc*, *Tacr1*, and both markers. (n = 8; *Tacr1* M = 18.25, SEM = 2.644; *Pkdcc* M = 41.00, SEM = 3.140, Double Positive M = 11.75, SEM = 2.016). **(C)** A breakdown of expression in B by sex shows no significant difference between males and females (n = 4 per sex; Two-way ANOVA F (1, 18) = 4.141, P=0.0569.) DP = double positive.

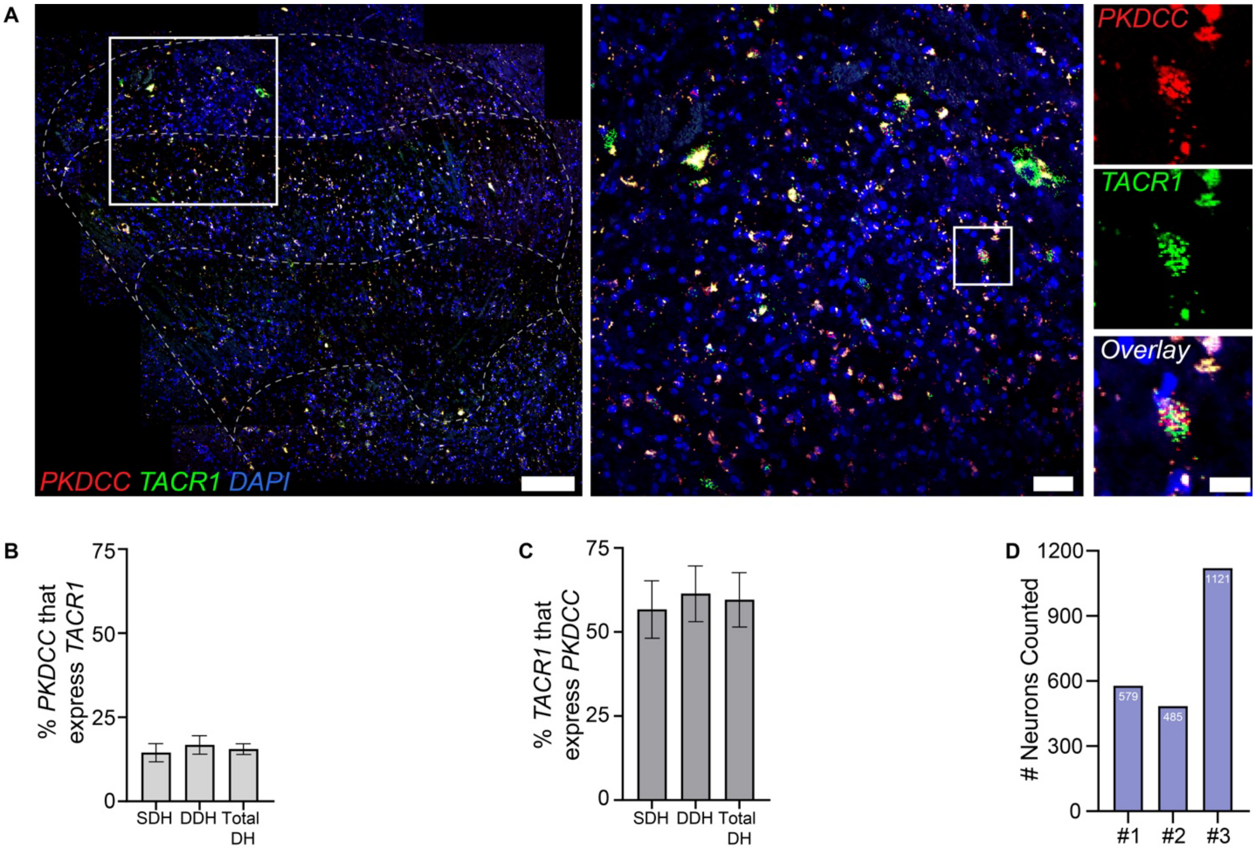


**Fig. S7. *PKDCC* is expressed in the human lumbar spinal cord dorsal horn.** (**A**) RNAscope *in situ* hybridization was used to identify *PKDCC* and TACR1-expressing neurons.Scale bar= 50 µm.Inset shows superficial dorsal horn from (A), the right panels display an example of a double positive neuron from the image. Scale bar= 5 µm (**B**) Percent of *PKDCC*+ neurons across the DH lamina that also express *TACR1*. (**C**) Percentage of *TACR1*$^{+}$ neurons that also express *PKDCC*. (**D**), a summary of total neurons counted for each donor (#1 – 3).

**Table S1. Donor information for tissues used in human characterization experiments.**

| Donor ID | Age | Sex | Ethnicity | Cause of Death | Experiment |
|---|---|---|---|---|---|
| 1 | 51 | M | White | CVA/Stroke | Fig 3 |
| 2 | 61 | F | White | Anoxia/Cardiac Arrest | Fig 3 |
| 3 | 19 | M | White | Anoxia/Drug Overdose | Fig 3 |
| 4 | 30 | M | Black | Anoxia/Asthma attack | Fig 3 |
| 5 | 34 | M | White | CVA/Stroke | Fig 3 |
| 6 | 34 | F | White | CVA/Stroke | Fig 3 |
| 7 | 41 | F | White | Head Trauma/Motorcycle Accident | Fig 3 |
| 8 | 36 | F | Black | Head Trauma/Blunt Injury/MVA | Fig 3 |
| 9 | 47 | M | White | CVA/Stroke | Fig S7 |
| 10 | 45 | F | White | Anoxia/Cardiac Arrest | Fig S7 |
| 11 | 19 | M | Hispanic | Head Trauma/GSW | Fig S7 |

**Table S2. Primers**

| Species | Gene | Primer |
|---|---|---|
| Human | *PKDCC* (VLK) | Fwd 5'-GAAGCGGAACCTCTATAATGCC-3' |
| Human | *PKDCC* (VLK) | Rev 5'-GGATACACTGGTACTCGGTACT-3' |
| Human | *DIPK1C* (Fam69C) | Fwd 5'-GAGGAGTACGTCTACTTCAGCC-3' |
| Human | *DIPK1C* (Fam69C) | Rev 5'-CGGTGGGAAAAGTCACTGTC-3' |
| Human | *DIPK2B* (Dia1R) | Fwd 5'-TGGACTGCCTCCCGATTCCT-3' |
| Human | *DIPK2B* (Dia1R) | Rev 5'- GGCTGATGCAGAGGCACAGA-3' |
| Human | *GAPDH* | Fwd 5'-ATGGGGAAGGTGAAGGTCG-3' |
| Human | *GAPDH* | Rev 5'-GGGGTCATTGATGGCAACAATA-3' |
| Mouse | *Pkdcc* (VLK) | Fwd 5'-CGGTCACCCTGCTTGACTTC-3' |
| Mouse | *Pkdcc* (VLK) | Rev 5'-GGCATTGTAGAGGTTCCGTTTC-3' |
| Mouse | *Dipk1c* (Fam69C) | Fwd 5'-GCCCTCAGCTTCCTGGACAT-3' |
| Mouse | *Dipk1c* (Fam69C) | Rev 5'-ACGTCAATAGCTACCACCGT-3' |
| Mouse | *Dipk2b* (Dia1R) | Fwd 5'-GCCAGCTTCTCTCCCTCCTT-3' |
| Mouse | *Dipk2b* (Dia1R) | Rev 5'-AGGTGGGAAGCCAGTGAGTT-3' |
| Mouse | *Hprt* | Fwd 5'- TCAGTCAACGGGGGACATAAA-3' |
| Mouse | *Hprt* | Rev 5'-GGGGCTGTACTGCTTAACCAG-3' |
| Mouse | *5' LoxP Pkdcc* | Fwd 5'- CTACAGATAATCTCCTTTAATGGGTGGG-3' |
| Mouse | *5' LoxP Pkdcc* | Rev 5'-CTTCTACGCAGAGTACTTCAGAAACC-3' |
| Mouse | *3' LoxP Pkdcc* | Fwd 5'- GACTCCTTGGTCTTACGGAG-3' |
| Mouse | *3' LoxP Pkdcc* | Rev 5'-CTCCACCAGCTTTAGCCAGC-3' |

**Movie S1. VLK is enriched at presynaptic terminals.**
Live cell imaging of axon segment from cultured rat cortical neurons (DIV21-23) transfected with VLK-mCherry (magenta) and Synaptophysin1-EGFP (green). The duration of imaging was 5 min. Colored arrowheads indicate puncta that are stationary for the duration of imaging. Pink and Yellow arrowheads are examples of stationary VLK localized with stationary Syp1 and Orange arrowhead stationary Syp1 alone (Also refer Fig. 3, B and C).

### References


1. V. S. Tagliabracci *et al.*, A Single Kinase Generates the Majority of the Secreted Phosphoproteome. *Cell* **161**, 1619-1632 (2015).
2. V. S. Tagliabracci *et al.*, Secreted kinase phosphorylates extracellular proteins that regulate biomineralization. *Science* **336**, 1150-1153 (2012).
3. Y. Imuta, N. Nishioka, H. Kiyonari, H. Sasaki, Short limbs, cleft palate, and delayed formation of flat proliferative chondrocytes in mice with targeted disruption of a putative protein kinase gene, Pkdcc (AW548124). *Dev Dyn* **238**, 210-222 (2009).
4. H. R. Washburn, N. L. Xia, W. Zhou, Y. T. Mao, M. B. Dalva, Positive surface charge of GluN1 N-terminus mediates the direct interaction with EphB2 and NMDAR mobility. *Nat Commun* **11**, 570 (2020).
5. H. R. Washburn, P. Chander, K. D. Srikanth, M. B. Dalva, Transsynaptic Signaling of Ephs in Synaptic Development, Plasticity, and Disease. *Neuroscience* **508**, 137-152 (2023).
6. K. Hanamura *et al.*, Extracellular phosphorylation of a receptor tyrosine kinase controls synaptic localization of NMDA receptors and regulates pathological pain. *PLoS Biol* **15**, e2002457 (2017).
7. M. J. Nolt *et al.*, EphB controls NMDA receptor function and synaptic targeting in a subunit-specific manner. *J Neurosci* **31**, 5353-5364 (2011).
8. M. S. Kayser, A. C. McClelland, E. G. Hughes, M. B. Dalva, Intracellular and trans-synaptic regulation of glutamatergic synaptogenesis by EphB receptors. *J Neurosci* **26**, 12152-12164 (2006).
9. M. B. Dalva *et al.*, EphB receptors interact with NMDA receptors and regulate excitatory synapse formation. *Cell* **103**, 945-956 (2000).
10. C. Minkin, Bone acid phosphatase: tartrate-resistant acid phosphatase as a marker of osteoclast function. *Calcif Tissue Int* **34**, 285-290 (1982).
11. H. Rico, L. F. Villa, Serum tartrate-resistant acid phosphatase (TRAP) as a biochemical marker of bone remodeling. *Calcif Tissue Int* **52**, 149-150 (1993).
12. M. J. Zylka *et al.*, Prostatic acid phosphatase is an ectonucleotidase and suppresses pain by generating adenosine. *Neuron* **60**, 111-122 (2008).
13. P. Guedes-Dias, E. L. F. Holzbaur, Axonal transport: Driving synaptic function. *Science* **366**, (2019).
14. N. Ahimsadasan, V. Reddy, M. Z. Khan Suheb, A. Kumar, in *StatPearls*. (StatPearls Publishing
Copyright © 2024, StatPearls Publishing LLC., Treasure Island (FL), 2024).
15. S. I. Sheffler-Collins, M. B. Dalva, EphBs: an integral link between synaptic function and synaptopathies. *Trends Neurosci* **35**, 293-304 (2012).
16. A. Y. Kim *et al.*, Pirt, a phosphoinositide-binding protein, functions as a regulatory subunit of TRPV1. *Cell* **133**, 475-485 (2008).
17. A. Latremoliere, C. J. Woolf, Central sensitization: a generator of pain hypersensitivity by central neural plasticity. *J Pain* **10**, 895-926 (2009).
18. S. N. Hassler *et al.*, The cellular basis of protease-activated receptor 2–evoked mechanical and affective pain. *JCI Insight* **5**, (2020).
19. E. V. Brown, A. F. Malik, E. R. Moese, A. F. McElroy, A. C. Lepore, Differential Activation of Pain Circuitry Neuron Populations in a Mouse Model of Spinal Cord Injury-Induced Neuropathic Pain. *J Neurosci* **42**, 3271-3289 (2022).
20. A. Barik *et al.*, A spinoparabrachial circuit defined by Tacr1 expression drives pain. *Elife* **10**, (2021).
21. S. Choi *et al.*, Parallel ascending spinal pathways for affective touch and pain. *Nature* **587**, 258-263 (2020).

22. A. M. Bell *et al.*, Deep sequencing of Phox2a nuclei reveals five classes of anterolateral system neurons. *bioRxiv*, (2023).
23. O. Gautier *et al.*, Challenges of profiling motor neuron transcriptomes from human spinal cord. *Neuron* **111**, 3739-3741 (2023).
24. J. A. Blum *et al.*, Single-cell transcriptomic analysis of the adult mouse spinal cord reveals molecular diversity of autonomic and skeletal motor neurons. *Nat Neurosci* **24**, 572-583 (2021).
25. T. L. Daigle *et al.*, A Suite of Transgenic Driver and Reporter Mouse Lines with Enhanced Brain-Cell-Type Targeting and Functionality. *Cell* **174**, 465-480.e422 (2018).
26. T. Huang *et al.*, Identifying the pathways required for coping behaviours associated with sustained pain. *Nature* **565**, 86-90 (2019).
27. B. S. Khakh, Molecular physiology of p2x receptors and atp signalling at synapses. *Nature Reviews Neuroscience* **2**, 165-174 (2001).
28. J. P. Dupuis, O. Nicole, L. Groc, NMDA receptor functions in health and disease: Old actor, new dimensions. *Neuron* **111**, 2312-2328 (2023).
29. R. C. Malenka, R. A. Nicoll, NMDA-receptor-dependent synaptic plasticity: multiple forms and mechanisms. *Trends Neurosci* **16**, 521-527 (1993).
30. B. C. Park, M. Reese, V. S. Tagliabracci, Thinking outside of the cell: Secreted protein kinases in bacteria, parasites, and mammals. *IUBMB Life* **71**, 749-759 (2019).
31. R. Ruscheweyh, O. Wilder-Smith, R. Drdla, X.-G. Liu, J. Sandkühler, Long-Term Potentiation in Spinal Nociceptive Pathways as a Novel Target for Pain Therapy. *Molecular Pain* **7**, 1744-8069-1747-1720 (2011).
32. C. N. Sang, NMDA-Receptor Antagonists in Neuropathic Pain: Experimental Methods to Clinical Trials. *Journal of Pain and Symptom Management* **19**, 21-25 (2000).
33. M. Hruska, N. Henderson, S. J. Le Marchand, H. Jafri, M. B. Dalva, Synaptic nanomodules underlie the organization and plasticity of spine synapses. *Nature Neuroscience* **21**, 671-682 (2018).
34. Y. T. Mao *et al.*, Filopodia Conduct Target Selection in Cortical Neurons Using Differences in Signal Kinetics of a Single Kinase. *Neuron* **98**, 767-782.e768 (2018).
35. M. R. Bordoli *et al.*, A secreted tyrosine kinase acts in the extracellular environment. *Cell* **158**, 1033-1044 (2014).
36. P. Chander, M. J. Kennedy, B. Winckler, J. P. Weick, Neuron-Specific Gene 2 (NSG2) Encodes an AMPA Receptor Interacting Protein That Modulates Excitatory Neurotransmission. *eneuro* **6**, ENEURO.0292-0218.2018 (2019).
37. D. Guez-Barber *et al.*, FACS purification of immunolabeled cell types from adult rat brain. *J Neurosci Methods* **203**, 10-18 (2012).
38. M. Hruska, N. T. Henderson, N. L. Xia, S. J. Le Marchand, M. B. Dalva, Anchoring and synaptic stability of PSD-95 is driven by ephrin-B3. *Nat Neurosci* **18**, 1594-1605 (2015).
39. A. J. B. Kreutzberger *et al.*, Reconstitution of calcium-mediated exocytosis of dense-core vesicles. *Science Advances* **3**, e1603208 (2017).
40. J. Rappsilber, M. Mann, Y. Ishihama, Protocol for micro-purification, enrichment, pre-fractionation and storage of peptides for proteomics using StageTips. *Nat Protoc* **2**, 1896-1906 (2007).
41. Y. Xu *et al.*, Cardiolipin remodeling enables protein crowding in the inner mitochondrial membrane. *Embo j* **40**, e108428 (2021).
42. J. Cox *et al.*, Andromeda: a peptide search engine integrated into the MaxQuant environment. *J Proteome Res* **10**, 1794-1805 (2011).
43. S. Tyanova, T. Temu, J. Cox, The MaxQuant computational platform for mass spectrometry-based shotgun proteomics. *Nat Protoc* **11**, 2301-2319 (2016).
44. R. Craig, J. P. Cortens, R. C. Beavis, Open source system for analyzing, validating, and storing protein identification data. *J Proteome Res* **3**, 1234-1242 (2004).

45. S. D. Sherrod *et al.*, Label-free quantitation of protein modifications by pseudo selected reaction monitoring with internal reference peptides. *J Proteome Res* **11**, 3467-3479 (2012).
46. S. Shiers, R. M. Klein, T. J. Price, Quantitative differences in neuronal subpopulations between mouse and human dorsal root ganglia demonstrated with RNAscope in situ hybridization. *PAIN* **161**, 2410-2424 (2020).
47. S. I. Shiers *et al.*, Convergence of peptidergic and non-peptidergic protein markers in the human dorsal root ganglion and spinal dorsal horn. *Journal of Comparative Neurology* **529**, 2771-2788 (2021).
48. S. R. Chaplan, F. W. Bach, J. W. Pogrel, J. M. Chung, T. L. Yaksh, Quantitative assessment of tactile allodynia in the rat paw. *J Neurosci Methods* **53**, 55-63 (1994).
49. D. J. Langford *et al.*, Coding of facial expressions of pain in the laboratory mouse. *Nature Methods* **7**, 447-449 (2010).
50. M. Cheah, J. W. Fawcett, M. R. Andrews, Assessment of Thermal Pain Sensation in Rats and Mice Using the Hargreaves Test. *Bio-protocol* **7**, e2506 (2017).
51. D. S. Brenner, J. P. Golden, R. W. t. Gereau, A novel behavioral assay for measuring cold sensation in mice. *PLoS One* **7**, e39765 (2012).
52. A. M. Cowie, C. L. Stucky, A Mouse Model of Postoperative Pain. *Bio-protocol* **9**, e3140 (2019).
